# The Integrated Cellular and Molecular Landscape of Autoimmunity

**DOI:** 10.1101/2025.09.12.675833

**Authors:** Romina Appierdo, Gerardo Pepe, Manuela Helmer-Citterich, Marina Sirota, Francesco Vallania, Pier Federico Gherardini

## Abstract

We performed a large-scale immunogenomic analysis of ∼13,000 transcriptomic profiles from 10 autoimmune diseases, integrating publicly available datasets from both blood and tissue. Using meta-analysis, we identified core immune mechanisms underlying autoimmunity, including strong interferon responses, inflammation, and adaptive immune suppression, alongside disease-specific signatures. To enhance biological interpretability, we derived higher-order immune features - such as cell type proportions, cytokine levels, pathway activity, transcription factor regulation, and miRNA activity - and organized them into 15 coherent immune modules. These modules enabled systematic cross-disease comparisons, revealing shared and distinct immunopathological patterns. The inflammation module, in particular, was associated with disease severity and predicted treatment response across multiple conditions. This modular framework offers a powerful tool for understanding immune dysregulation and advancing precision medicine in autoimmune diseases. To support reproducibility and enable others to build upon this work, we developed an interactive app to explore and download the complete dataset and associated results.

## Introduction

Autoimmune diseases encompass a diverse group of disorders characterized by the immune system mistakenly targeting the body’s own tissues. This misdirected immune response leads to chronic inflammation and tissue damage, with specific conditions affecting nearly every organ system.^1,2^ Despite their significant impact on global health, with an estimated increasing prevalence of ∼12% annually,^3^ autoimmune diseases remain poorly understood.

The complexity of autoimmune diseases is driven by the intricate interplay between genetic predisposition and environmental triggers.^1^ Many of them share common genetic risk loci, particularly within the major histocompatibility complex (MHC), which plays a central role in immune regulation.^4^ However, genetic susceptibility alone does not fully explain disease onset. Environmental exposures act as key triggers that influence disease development and progression.^1,5^ Factors such as ultraviolet (UV) radiation, viral infections, and lifestyle changes contribute to immune dysregulation. For example, Epstein-Barr virus (EBV) infection has been implicated in multiple autoimmune conditions,^6,7^ while smoking and gut microbiome alterations are known to influence disease risk and severity, such as systemic lupus erythematosus (SLE) and inflammatory bowel diseases (IBDs).^8,9^

The heterogeneity in the etiology and clinical manifestations of autoimmune diseases presents a major challenge for both research and treatment development.^10,11^ Conditions such as type 1 diabetes (T1D) and Hashimoto’s thyroiditis primarily affect specific organs, while others, like SLE and rheumatoid arthritis (RA), involve multiple systems. This variability complicates disease classification, diagnosis, and the identification of shared underlying mechanisms.

While substantial progress has been made in the characterization of individual autoimmune diseases, a systematic, data-driven comparative survey of their immune features remains lacking. Similar large-scale efforts have been transformative in oncology, where pan-cancer immune landscape studies have revealed common biological pathways and therapeutic targets across different malignancies.^12^ A comparable approach in autoimmunity could uncover shared immune mechanisms that drive disease pathogenesis, identify disease-specific processes, and inform new therapeutic strategies.

To address this gap, we leveraged the growing body of publicly available transcriptomic data to characterize and dissect the immune landscape of autoimmunity. We used a meta-analysis framework to integrate data from 10 autoimmune diseases, compiling 6,238 blood samples from 53 studies and 7,025 tissue samples from 103 studies across five tissue types. Data integration of multiple independent studies offers key advantages in this context: it enhances statistical power, mitigates study-specific biases, and enables robust cross-disease comparisons.

To enhance interpretability, we extracted higher-order features from transcriptomic data, including cytokine expression, immune cell abundance, pathway activity, transcription factor (TF) regulation, and miRNA activity. Using meta-analysis, we identified robust immune signatures across diseases in both blood and tissue, enabling large-scale comparisons. By further combining these features, we defined 15 modules representing key immune processes such as interferon (IFN) responses, TLR/NF-κB signaling, and adaptive immune suppression. Finally, we validated their biological and clinical relevance using independent datasets, including perturbation experiments and clinical cohorts.

By integrating large-scale transcriptomic datasets across diseases and tissue types, our study provides a framework for identifying both shared and disease-specific mechanisms of immune dysregulation. Our approach advances our understanding of autoimmune pathobiology guiding the development of novel therapeutic strategies.

## Results

### Overview of the workflow

We developed a computational pipeline to systematically characterize immune dysregulation across diseases by integrating public transcriptomes from a broad range of clinical studies. We chose transcriptomics data to derive higher order biologically interpretable features, given its ability to intrinsically capture both genetic and environmental factors, two key drivers of autoimmune disorders (**Figure 1**).

**Figure 1.**
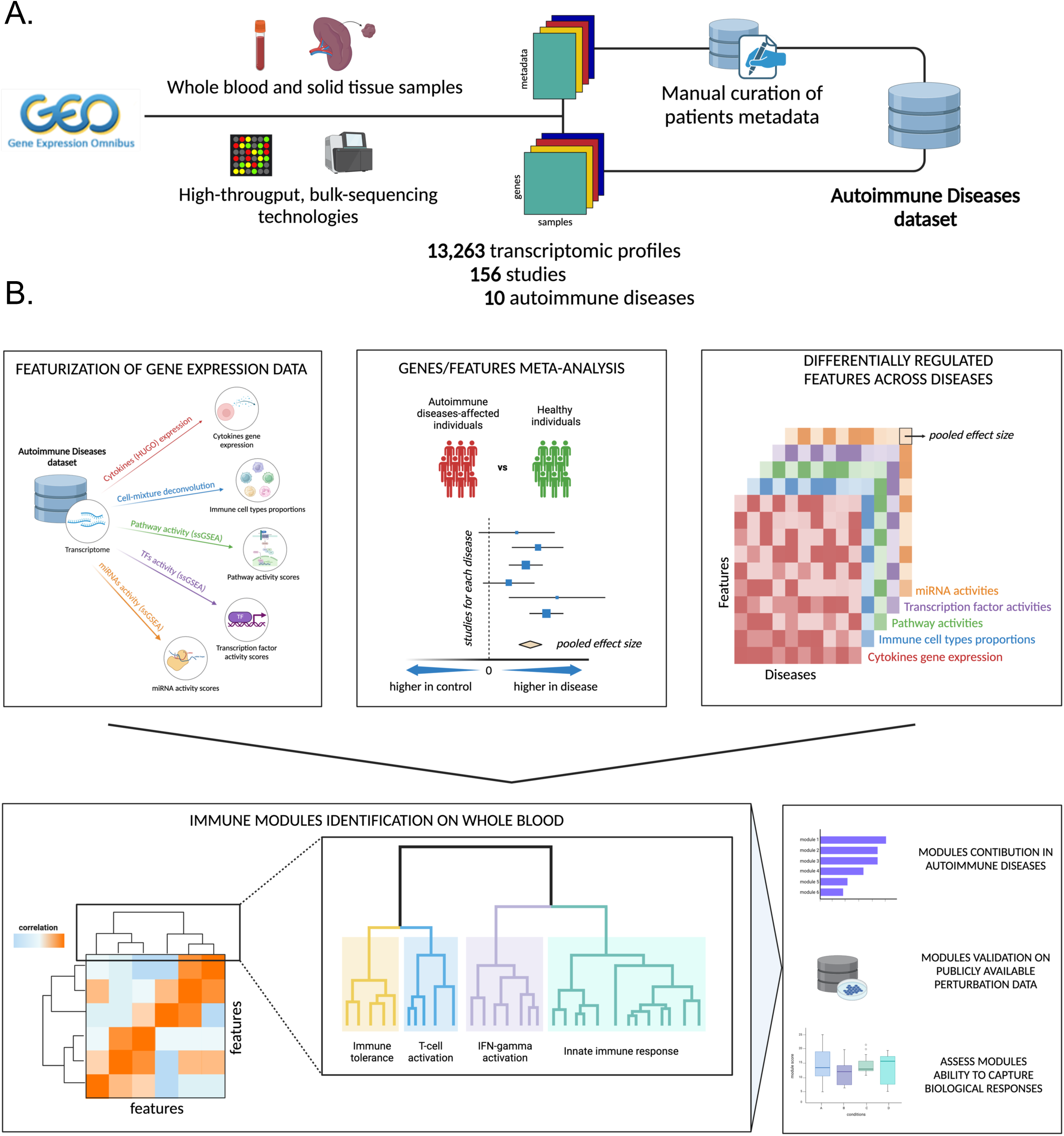
Project Workflow. **(A)** Collection and curation of gene expression data and metadata from the GEO database to create an Autoimmune Diseases dataset. **(B)** Featurization of gene expression data to obtain higher-order features, followed by meta-analysis at the gene and feature levels to identify a landscape of differentially regulated features across all diseases (top). Modules were identified and assigned a biological interpretation, followed by validation on autoimmune disease datasets, public perturbation data, and assessment of modules’ ability to capture biological responses and predict clinical outcomes (bottom). Figure created with BioRender.com

We collected, processed, and manually annotated transcriptomic data for 13,263 samples from the Gene Expression Omnibus (GEO) database,^13,14^ comprising blood and tissue samples for 10 autoimmune diseases, including systemic lupus erythematosus (SLE), psoriasis (Ps), Crohn’s disease (CD), ulcerative colitis (UC), Sjögren’s syndrome (SS), rheumatoid arthritis (RA), systemic sclerosis (or scleroderma, SSc), type 1 diabetes (T1D), multiple sclerosis (MS), and juvenile idiopathic arthritis (JIA) (**Figure 1A**).

We extracted higher-order biologically interpretable features, reflective of immune system activity, from gene expression data, to provide functional insights into the pathobiology of autoimmune disorders (**Figure 1B, top-left panel**). These features encompass immune cell abundance, cytokine expression, pathway activity, TF regulation, and miRNA activity.

We next performed a meta-analysis to identify genes and immune features that were consistently dysregulated in autoimmune diseases across multiple studies, revealing robust disease-specific and pan-disease signatures (**Figure 1B, top-middle and right panels**).

We then developed a computational pipeline to identify groups of highly co-regulated features across diseases (**Figure 1B, bottom-left panel**). Features clustering across autoimmune diseases identified 15 biologically meaningful modules representing key immunological mechanisms, including IFN signaling, TLR/NF-κB activation, and adaptive immune triggering. These modules provided a structured framework for describing immune dysfunction. We leveraged independent datasets profiling perturbation experiments or clinical studies (**Figure 1B, bottom-right panel**) to validate their biological and clinical relevance.

Finally, we applied our modules to clinical datasets profiling patients before and after treatment, evaluating associations of specific modules - such as the inflammation module - with disease severity and predicted therapeutic outcomes.

### Peripheral Blood Gene Expression Meta-Analysis Highlights IFN Activation and Adaptive Immune Suppression Across Autoimmune Diseases

We integrated gene expression data from peripheral blood to identify robust disease-specific and pan-disease signatures. We analyzed whole blood transcriptomic data from all profiled autoimmune diseases, which comprises 6,238 samples including disease-affected and control individuals, across 53 studies (**Table 1**). To derive robust transcriptional signatures, we performed a meta-analysis of gene expression data across studies for each disease to standardize and pool differential expression changes between diseased and healthy individuals, averaging out dataset-specific confounders.^15^

**Table 1.**
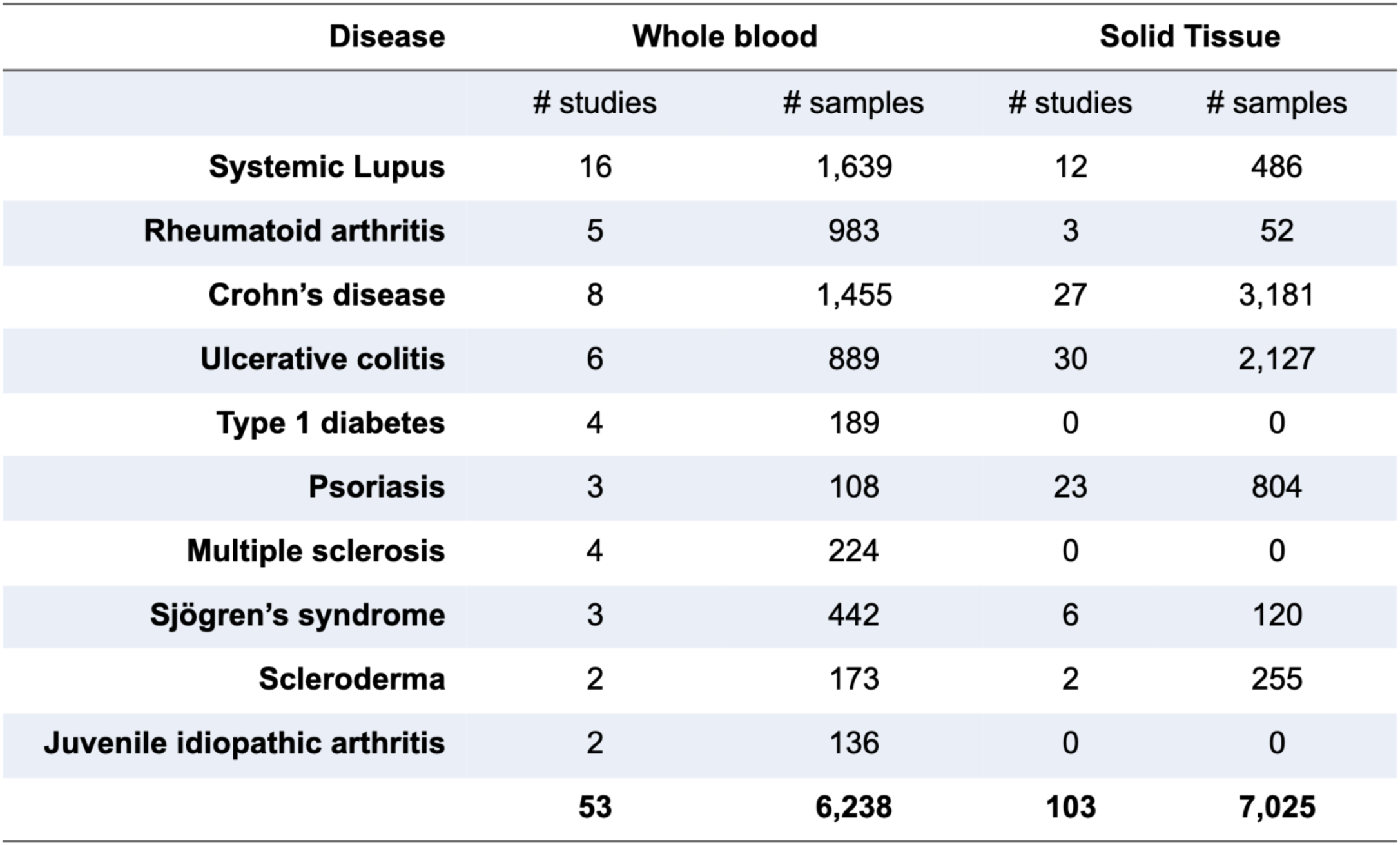
Summary of the curated dataset used for the analysis. The table reports the number of transcriptomic studies and samples included for each autoimmune disease, separated by compartment: whole blood and solid tissue. A total of 53 blood studies (6,238 samples) and 103 tissue studies (7,025 samples) were compiled across 10 autoimmune diseases. This manually curated dataset integrates publicly available data from GEO and forms the foundation for the cross-disease analyses presented in this study.

| Disease | Whole blood |  | Solid Tissue |  |
| --- | --- | --- | --- | --- |
|  | # studies | # samples | # studies | # samples |
| Systemic Lupus | 16 | 1,639 | 12 | 486 |
| Rheumatoid arthritis | 5 | 983 | 3 | 52 |
| Crohn's disease | 8 | 1,455 | 27 | 3,181 |
| Ulcerative colitis | 6 | 889 | 30 | 2,127 |
| Type 1 diabetes | 4 | 189 | 0 | 0 |
| Psoriasis | 3 | 108 | 23 | 804 |
| Multiple sclerosis | 4 | 224 | 0 | 0 |
| Sjögren's syndrome | 3 | 442 | 6 | 120 |
| Scleroderma | 2 | 173 | 2 | 255 |
| Juvenile idiopathic arthritis | 2 | 136 | 0 | 0 |
|  | 53 | 6,238 | 103 | 7,025 |

We compared aggregated gene expression changes across autoimmune conditions, focusing on genes that were significantly up- or down-regulated (FDR<0.05) in at least one disease (**Supplementary File 1**). Hierarchical clustering of diseases using only significant genes (**Figure 2A**) revealed similar (ρ=0.88, p<2.2e-16) expression patterns between CD and UC, reflecting their shared molecular and clinical features as IBDs.^16^ A separate cluster included RA, SSc, SLE, and SS, all of which are rheumatic diseases.^17,18^ We observed strikingly similar transcriptional changes across these diseases (**Supplementary Table 1**), reflecting the known consistent upregulation of shared pathways, including IFN signaling and inflammatory processes.^17,19,20^

**Figure 2.**
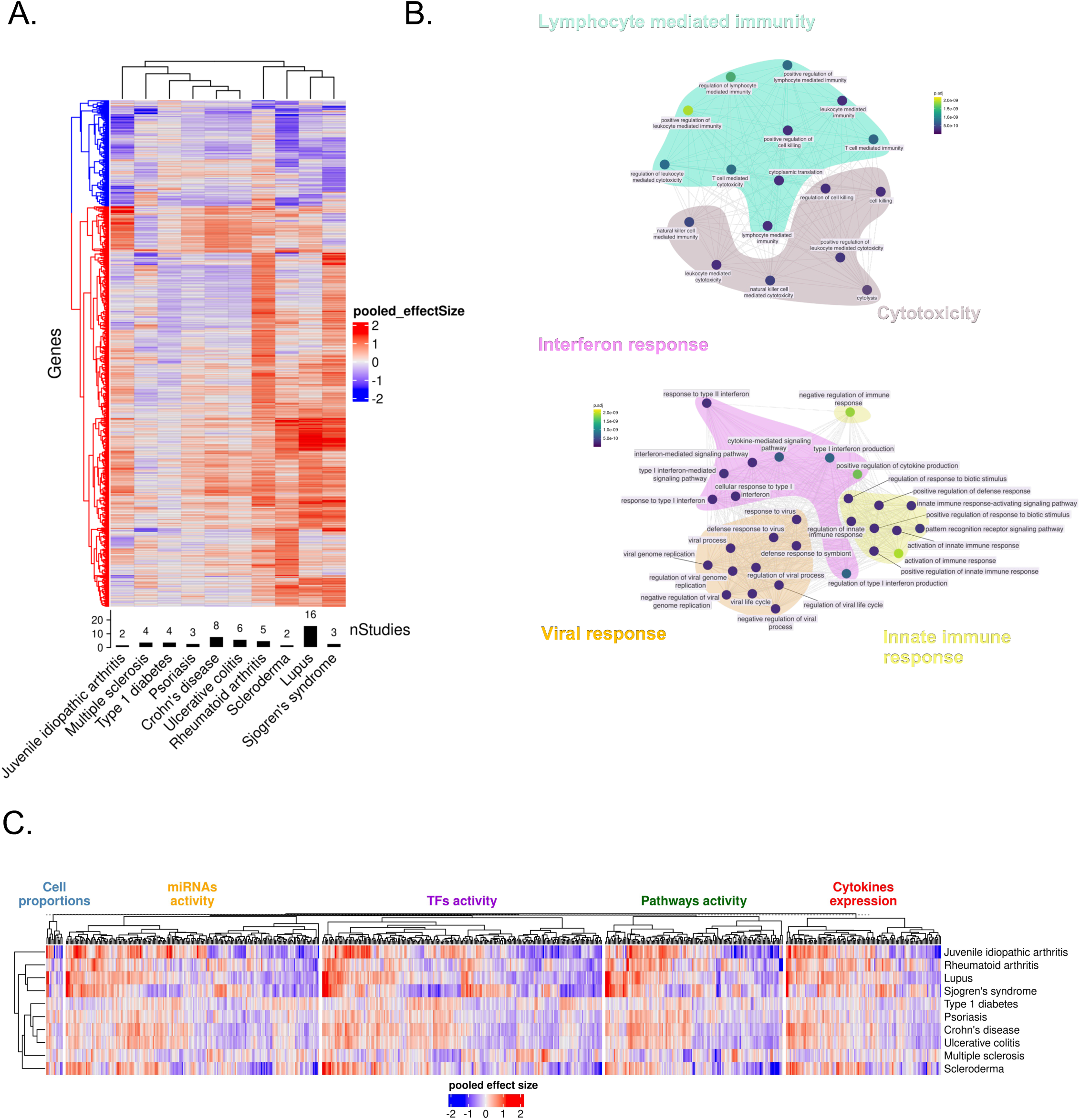
Resolving the Multi-Layered Immune Landscape of Autoimmune Diseases Through Gene Expression Meta-Analysis and Feature Integration. **(A)** Gene expression meta-analysis on whole blood samples. Rows of the heatmap represent genes; only genes with a significant effect size in at least one disease are shown. Color indicates the pooled effect size, with red representing upregulation and blue representing downregulation. Hierarchical clustering identifies two gene clusters: one with genes mostly upregulated across diseases (red cluster) and one with genes mostly downregulated (blue cluster). **(B)** Enrichment analysis on the upregulated gene cluster (bottom). Enrichment analysis on the downregulated gene cluster (top). **(C)** Landscape of deregulated features across diseases. Columns represent features that are significant in at least one disease. Red indicates upregulation, blue indicates downregulation. Rows show disease clustering based on all features.

We also identified a separate cluster composed of more heterogeneous diseases, including IBDs, MS, JIA, T1D and Ps. Despite exhibiting distinct expression patterns, these diseases share common biological mechanisms.^21^ Similarities in genetic risk factors, autoimmune reactivity and genetic epidemiology between MS and T1D have been previously reported,^22–24^ and shared susceptibility loci have been identified between MS and IBDs.^21,25^

Clustering analysis (**Figure 2A**) revealed two distinct sets with either up- or downregulated genes across different disease groups. Genes in the upregulated cluster were enriched (FDR<0.05) in biological processes related to IFN responses, inflammation, innate immune responses, and viral responses (**Figure 2B, bottom**), all of which are well established mechanisms in autoimmunity.^26–28^ In contrast, the downregulated cluster was enriched for lymphocyte activation and cytotoxicity (**Figure 2B, top**), suggesting a deregulation of adaptive immune processes, including T cell receptor signaling and CD8+ T cell cytolytic activity.^29^ This aligns with previous observations that chronic inflammation in autoimmunity can lead to T cell exhaustion and impaired cytotoxic function.^30,31^

Together, our findings reveal convergent immune mechanisms - most notably, the upregulation of innate immune responses such as IFN signaling and inflammation, alongside the downregulation of adaptive immune components - that could serve as shared therapeutic targets.

### Deriving Higher-Order Features to Uncover the Biology of Autoimmune Diseases

To facilitate biological interpretation, we employed several computational approaches to extract higher-order features from the raw gene expression data.

We defined a set of five immune-related features - immune cell proportions, pathway, TF and miRNA activity scores, and cytokine levels - that captured complementary aspects of the immune system and enabled direct comparisons between samples.

We then developed a computational pipeline to calculate feature scores from gene expression data across all samples and datasets. By aggregating these features across multiple studies, we captured reproducible patterns of immune dysregulation that reflect both shared and disease-specific processes (**Supplementary File 1**).

We next asked whether featurization improved interpretability by revealing known immunological changes observed in specific autoimmune diseases. Our observed differences in cell proportions recapitulated previous observations, including lymphopenia in SLE (pooled_effect_size=-0.38, p=5.19e-09),^32,33^ and an overall increase in pro-inflammatory M1 macrophages (pooled_effect_size=0.41, p=5.2e-03)^34^ (**Supplementary Figure 1**).

Furthermore, we identified upregulated IFN signaling pathways (**Supplementary Figure 2** and **Supplementary File 1**), in line with its well-documented dysregulation in autoimmune diseases.^35^ Similarly, TF analysis revealed increased activity of key regulators, including members of the STAT and IRF families (**Supplementary Figure 3** and **Supplementary File 1**), which orchestrate IFN-mediated immune responses.^36^

Disease clustering with these features (**Figure 2C**) broadly recapitulated the same structure obtained using gene expression. Consistently, IBDs clustered together, whereas SS and SLE grouped closely, reflecting their shared pathogenic mechanisms.^17^

Notably, feature-based clustering provided greater resolution of disease relationships, such as that between SLE and RA - an association extensively supported by the literature.^37,38^ Overall, our results suggest that feature-level analyses capture key immunological processes with greater clarity, enhancing the ability to identify cross-disease connections in autoimmunity.

### Tissue Analysis Captures Localized and Disease-Specific Immune Dysregulation

Peripheral blood is widely used due to its minimally invasive collection, but fails to recapitulate tissue immunology and, consequently, core disease mechanisms.^39^ Therefore, we extended our analysis to 7,025 transcriptomes from samples of the primary affected tissue across 7 autoimmune diseases (**Table 1**), including kidney and skin for SLE, synovium for RA, salivary glands for SS, skin for Ps and SSc, and gut for UC and CD. We performed gene expression meta-analysis on the tissue datasets (**Supplementary File 1**). Gene expression clustering across tissues revealed two distinct groups of genes exhibiting either systemic up- or down-regulation across most diseases (**Supplementary Figure 4A**). The upregulated cluster was significantly (FDR<0.05) enriched in (**Supplementary Figure 4B, bottom**) adaptive immunity, IFN responses, viral defense mechanisms, and cytokine signaling pathways, reflecting strong inflammatory activity in affected tissues, consistent with heightened immune activation typically observed at disease sites.^40^ In contrast, the downregulated cluster showed enrichment (FDR<0.05) for tissue structure and extracellular matrix (ECM) remodeling (**Supplementary Figure 4B, top**), reflecting the detrimental impact of autoimmune processes on tissue integrity and repair.^41–43^

We observed numerous molecular associations across different diseases: notably, lupus nephritis (LN) showed a distinct differential expression pattern, with a large proportion of genes upregulated in affected kidney tissue. Focusing on significantly elevated genes in LN (1,662 genes, FDR<0.05), we observed strong activation of IFN-α (fold_enrichment=4.5, FDR=3.3e-11) and γ (fold_enrichment=3.5, FDR=1.9e-13) signaling (**Supplementary Figure 5**), consistent with the well-documented role of IFN responses in SLE pathogenesis.^17,19,20,35,44^ We also identified significant enrichment in complement activation (fold_enrichment=1.8, FDR=3.8e-02), a critical driver of LN, as the underlying immune dysfunction in this condition stems from the deposition of autoantibodies and immune complexes, which initiate kidney inflammation through complement system activation.^41,45^

By comparing disease-specific gene expression effect sizes across disease pairs (**Supplementary Table 2**) we observed a strong correlation between SLE and Ps skin (ρ=0.74, p<2.2e-16). Molecular similarities between these conditions are unlikely to be driven by comorbidity, as SLE and Ps rarely co-occur.^46,47^ Rather, this suggests shared immunopathological mechanisms. Indeed, both diseases present dysregulated IFN signaling, heightened activation of plasmacytoid dendritic cells (pDCs),^48,49^ and an inflammatory milieu enriched in type I IFN-stimulated genes.^50^ Additionally, skin involvement in SLE, such as cutaneous lupus erythematosus, shares histopathological features with psoriatic inflammation, including keratinocyte hyperproliferation,^51,52^ and immune cell infiltration.^53,54^

We next compared Ps and RA and observed a moderate but significant correlation (ρ=0.28, p<2.2e-16). Interestingly, this correlation increased substantially (ρ=0.62, p<2.2e-16) when restricted to consider genes significant in both conditions, consistent with the presence of a shared molecular component despite the distinct clinical manifestations (Ps affecting skin; RA affecting joints). Psoriatic arthritis, a pathological complication of Ps that shares clinical and immunological features with both diseases^55^, may partially account for shared molecular processes. The involvement of shared immune pathways (e.g., TLR/NF-κB, IL-17, TNF-α) and Th17-driven inflammation supports this connection.^55^

Finally, we observed similarities between skin in SSc and salivary glands in SS (ρ=0.38, p=2.6e-4), two diseases that affect distinct tissues but share key molecular and pathogenic features.^56^ Both conditions are marked by chronic inflammation and dysregulated IFN responses, suggesting that common therapeutic strategies may benefit both diseases.^57^

Overall, large-scale integration of tissue-specific data revealed shared inflammatory mechanisms and molecular similarities across diseases.

### Tissue-Specific Immune Signatures Show Limited Concordance with Blood Profiles in Autoimmune Diseases

Autoimmune diseases involve both systemic and localized immune responses, but the extent of their similarity remains unclear.

To this end, we first quantified the number of significant features shared or unique between blood and tissue for each feature set (**Figure 3A**). Within each disease, the proportion of shared features was generally low, typically ranging from ∼5% to ∼20%, broadly consistent across different feature types.

**Figure 3.**
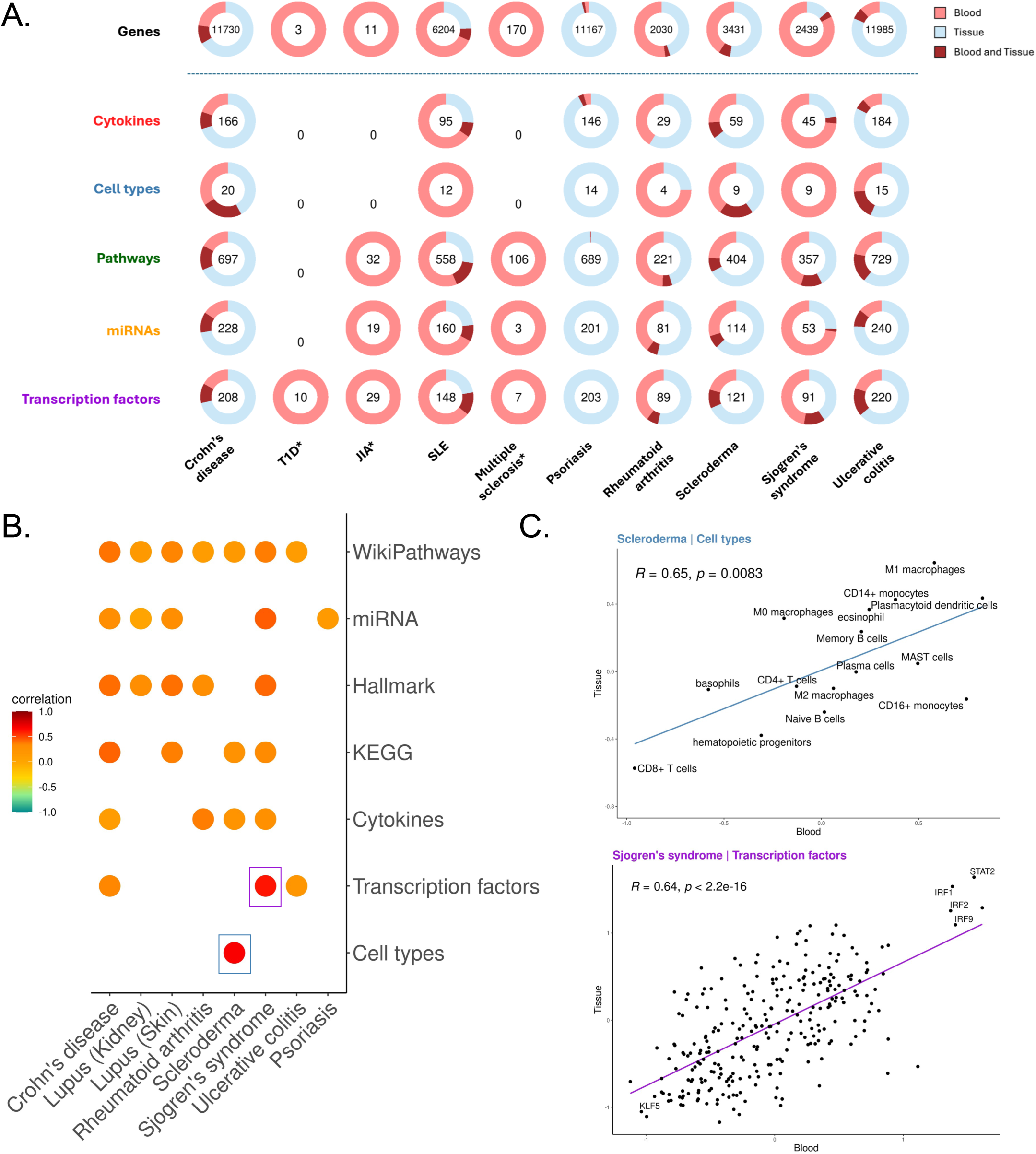
Comparative Analysis of Transcriptomics and Feature Data Reveals Both Distinct Differences and Shared Characteristics Between Tissue and Peripheral Immune Signatures. **(A)** Qualitative assessment of shared deregulated genes and features between blood and solid tissue for each disease. Each pie chart shows the number of significant deregulated features in both compartments, with the number in the center representing the union of significant features across blood and tissue (dark red). Light red indicates features significant only in blood, light blue only in tissue, and dark red in both blood and tissue. An asterisk (*) indicates diseases where only blood samples are available. **(B)** Spearman correlations of effect sizes in blood vs. tissue for each disease. Only correlations with p<0.05 are shown. Highest correlation values are highlighted with a box. **(C)** Close-up of correlation plots for transcription factors in SS and cell types in SSc, showing strong positive correlations.

The proportion of shared features between blood and tissue was high for several diseases (e.g. CD and UC, mean_fraction_of_overlapping_features=0.204 and 0.174, respectively), while others exhibited distinct compartment-specificity (e.g. RA and SS, mean_fraction_of_overlapping_features=0.041 and 0.073, respectively) (**Figure 3A**). These findings suggest that similarities between peripheral blood and tissue are disease-dependent.

We next assessed concordance of gene expression changes between blood and tissue (**Supplementary Figure 6**). Notably, we found no consistent pattern of coordinated regulation between blood and tissue (mean_Spearman_correlation_across_diseases, ρ=0.33), suggesting distinct regulatory mechanisms or compartment-specific immune responses within each disease.

We next quantified blood-tissue concordance of our immunological feature sets across diseases. Overall, most feature types showed only moderate correlation (ρ_median_=0.3) (**Figure 3B**), indicating low systemic concordance with tissue-specific immune activity. However, we observed strong positive correlations in several feature-disease pairs, such as cell-type abundance in SSc (ρ=0.68, p=8e-4) (**Figure 3C, top**) and TF activity in SS (ρ=0.62, p=4.95e-30) (**Figure 3C, bottom**). While broad immune dysregulation in blood does not consistently mirror tissue immunology, our results suggest that select features display cross-compartment consistency, highlighting potential systemic biomarkers for disease-specific immune activity.

Previous studies have established a strong link between disease-specific immune alterations in peripheral blood and corresponding changes in tissue-infiltrating immune cells. Building on this, we investigated the relationship between immune cells in periphery and tissue to better understand their interplay in autoimmune diseases.

Our analysis revealed distinct patterns of immune cell enrichment across diseases. Among all immune subsets, M1 macrophages exhibited a consistent trend of upregulation in both blood and tissue (mean_effect_size_across_diseases=0.39 and 0.75, respectively) (**Supplementary Figures 1 and 8**). These pro-inflammatory macrophages were significantly elevated in blood (median_effect_size_across_diseases=0.347) and tissue (median_effect_size_across_diseases=0.555). Their enrichment suggests a key role in sustaining inflammation, as they secrete cytokines such as TNF-α, IL-6, and IL-1β, which are central mediators of chronic immune activation in autoimmunity.^34^

In contrast, M0 macrophages (undifferentiated precursors) and M2 macrophages were not consistently enriched across all diseases (**Supplementary Figures 1 and 8 and Supplementary File 1**), likely reflecting differences between inflammatory and reparative processes in different autoimmune conditions.^34,58^

Our findings emphasize a consistent role of M1 macrophages in autoimmune inflammation, bridging systemic and localized immune responses. These results provide insights into macrophage-driven mechanisms in autoimmunity and suggest that targeting M1 macrophages could be a potential strategy for managing inflammation in multiple diseases.

Finally, we observed a consistent upregulation of pDCs (**Supplementary Figures 1 and 8**) in both tissue and periphery across all diseases (mean_effect_size_across_diseases=0.57 and 0.66, respectively). As primary producers of IFN, pDCs play a crucial role in driving immune dysregulation and inflammation in autoimmunity,^59,60^ further underscoring the strong connection between systemic and localized immune responses.

Our findings demonstrate that while peripheral immune activity does not reliably mirror tissue pathology in autoimmune diseases, several specific immune features - such as IFN signaling, M1 macrophage enrichment, and pDC activation - were concordant across compartments. These results highlight both the limitations of using peripheral blood as a proxy for tissue-level immune processes and the potential of specific immune signatures as systemic biomarkers.

### IFN Responses and Plasmacytoid Dendritic Cells as a Unifying Link Between Blood and Tissue in Autoimmune Diseases

Given the well-established role of IFNs in autoimmune pathogenesis, we systematically evaluated IFN activity across compartments and diseases.

IFN signaling plays a central role in the immune dysregulation underlying autoimmune diseases.^61^ By analyzing both blood and tissue samples, we observed consistent upregulation of IFN-related genes and pathways across multiple diseases, with enrichment in both type I and II IFN responses (**Supplementary Figure 7A** and **Supplementary File 1**). Notably, SLE displayed the strongest IFN signal (mean effect sizes in blood, kidney and skin were, respectively, 0.61, 1.02 and 1.14), in line with its recognition as a prototypical IFN-driven disease.^44^

We asked whether upregulation of the IFN signaling pathway would be biologically reflected throughout our different feature sets. Meta-analysis of immune cell proportions in both blood (**Supplementary Figure 7B** and **Supplementary File 1**) and tissue (**Supplementary Figure 7C** and **Supplementary File 1**) revealed a marked increase in pDCs across nearly all diseases, with the strongest signals seen in SLE (blood: effect_size=1.13, FDR=1.46e-08 ; skin: effect_size=1.72, FDR=0.046), SSc (blood: effect_size=0.82, FDR=0.02 ; skin: effect_size=0.43, FDR=0.02), and SS (blood: effect_size=0.83, FDR=7.5e-08). pDCs are the primary source of IFN,^60^ and their activation is a hallmark of SLE.^62^ Expectedly, the IFN signaling pathway also exhibited the highest enrichment in tissues, across all diseases, compared to other pathways (“TYPE_II_INTERFERON_SIGNALING_IFNG” : mean_effect_size=1.94, Fisher’s combined p=3.7e-60) (**Supplementary Figure 7D**).

Together, these findings position IFN signaling as a core immunological process in autoimmunity, which is consistently activated across diseases and compartments, and underscore its relevance as a mechanistic bridge between systemic and tissue-level immune responses.

### Clustering of Features Identifies 15 Immune Modules Recapitulating the Biology of Autoimmune Diseases

Although our analyses provided a comprehensive view of immune dysregulation in autoimmunity, the sheer number of significant features hindered biological interpretation (**Figures 2C** and **Supplementary Figure 4C**). We therefore leveraged correlation patterns among features across autoimmune diseases to identify a compact set of biologically meaningful immune modules. We performed this analysis using blood data to minimize confounders derived from tissue-specific gene expression patterns.

We identified 700 features significant in at least one disease (FDR<0.05), including pathways, immune cell populations, TF, and cytokines (**Figure 2C**). To investigate the underlying structure of our features, we computed pairwise Spearman correlations between features across diseases and clustered the resulting correlation matrix to identify groups of highly co-regulated features (**Figure 4A**). Hierarchical clustering yielded 15 distinct feature groups (**Supplementary File 2**) (cluster number robustness significance by perturbation analysis: p<0.05).

**Figure 4.**
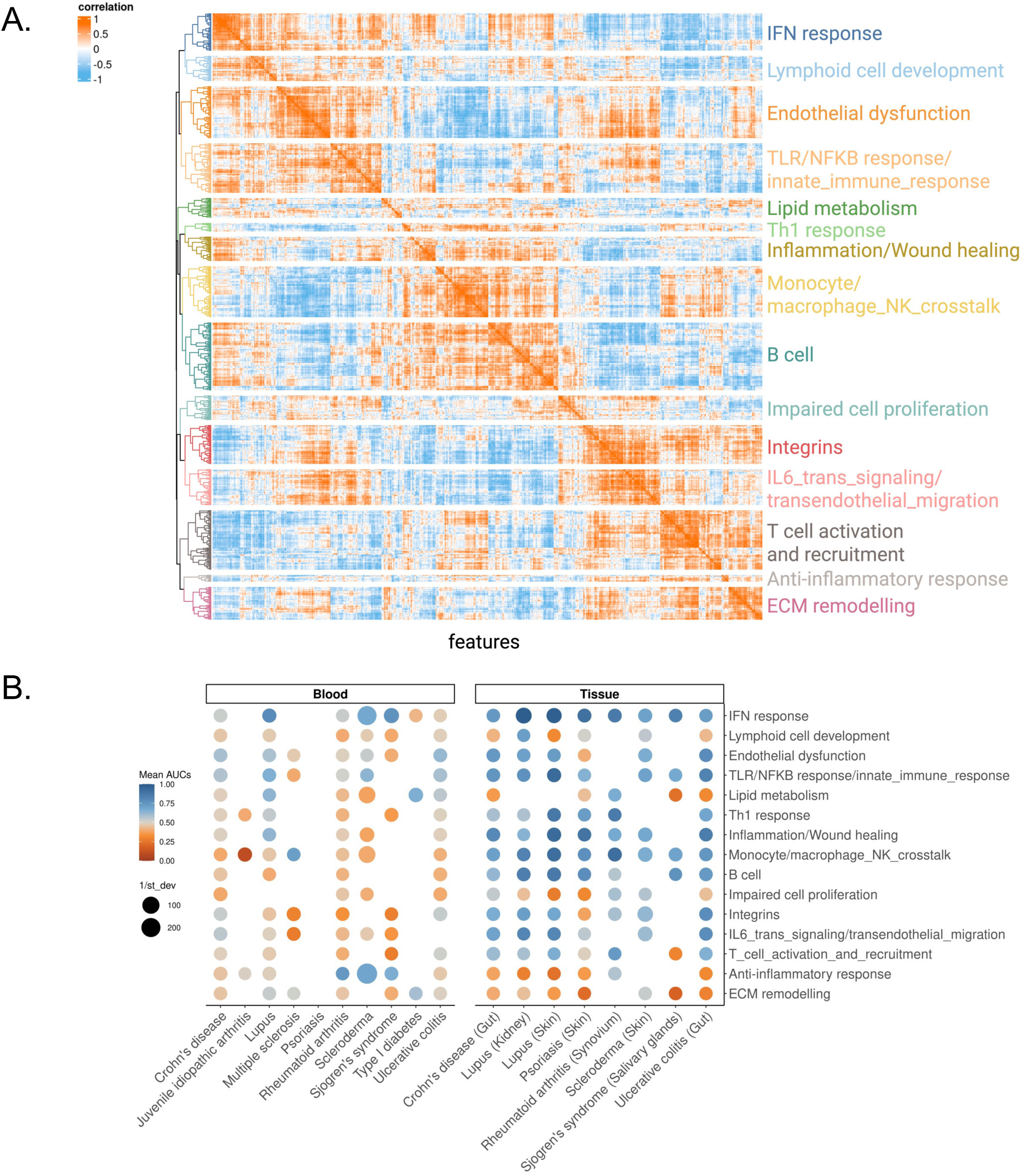
Identification of 15 Autoimmune-Related Modules and Their Contribution Across Diseases in Blood and Tissue. **(A)** Correlogram for module identification. Rows and columns represent 700 features that define the blood landscape. Spearman correlations were computed across all pairs of features, with orange indicating positive correlation and sky blue indicating negative correlation. Clustering was applied to identify groups of related features, resulting in 15 modules. Each module was labeled based on its biological significance. **(B)** Dot plot showing the contribution of each module to each disease across blood and tissue. Each dot represents the mean AUC, where AUCs were computed for each dataset per disease based on module activity and disease state labels. Blue indicates a high ability of the module to distinguish disease from control. Dot size represents 1/standard deviation, with larger dots indicating less heterogeneity across studies. Only significant (p<0.05) dots are shown.

We then manually annotated each cluster based on its constituent features, resulting in 15 immune modules that represented key immunological pathways and cellular programs (**Figure 4A**). For example, we identified modules corresponding to IFN responses, endothelial dysfunction, TLR/NF-κB signaling, and lipid metabolism. Other modules highlighted cellular processes, such as Lymphoid_cell_development, Monocyte/macrophage_NK_cell_crosstalk, and B cell activity. Our modules also included pathways related to impaired cell proliferation, ECM remodeling, and inflammation or wound healing, underscoring the diverse mechanisms driving these diseases.

### Immune Modules Capture IFN Response in Perturbation Dataset

To assess whether our modules truly captured underlying biological processes, we tested their behavior in controlled experimental settings using publicly available perturbation datasets.

Given the central role of IFNs in autoimmunity,^17,19,20,35,44,61^ we focused on the IFN response module by searching for transcriptomic datasets where the IFN pathway was experimentally perturbed. We selected a dataset profiling 24 whole blood samples from 8 healthy donors (GSE78193). Each sample was either stimulated with IFN-γ or left unstimulated, and gene expression was measured at three time points: baseline (unstimulated), 24 hours post-stimulation, and 48 hours post-stimulation.

To evaluate the IFN module responsiveness to stimulation, we assessed its activity across time points using both hierarchical clustering and quantitative scoring. At baseline, IFN-stimulated and unstimulated samples did not separate, indicating minimal pathway activation. However, at 24 and 48 hours, stimulated samples showed clear separation from unstimulated ones, with the strongest difference observed at 48 hours, likely reflecting the cumulative effect of prolonged IFN-γ exposure (**Figure 5A**). To quantify this response, we computed an aggregated module score per sample by summing activities across all IFN module features. Consistently, there was no significant difference in scores at baseline (Wilcoxon-test, p>0.05), while stimulated samples had significantly higher scores at 24 and 48 hours (Wilcoxon-test, p<0.05; **Figure 5B**). Together, these findings demonstrated that the IFN module robustly captures the biological response to IFN stimulation over time. To further validate that the observed differential activity of the IFN module was not due to random variation, we performed a permutation test (see Methods). The IFN module had a significantly higher effect size than expected by chance (p<0.001 for all time points; **Supplementary Figure 9**).

**Figure 5.**
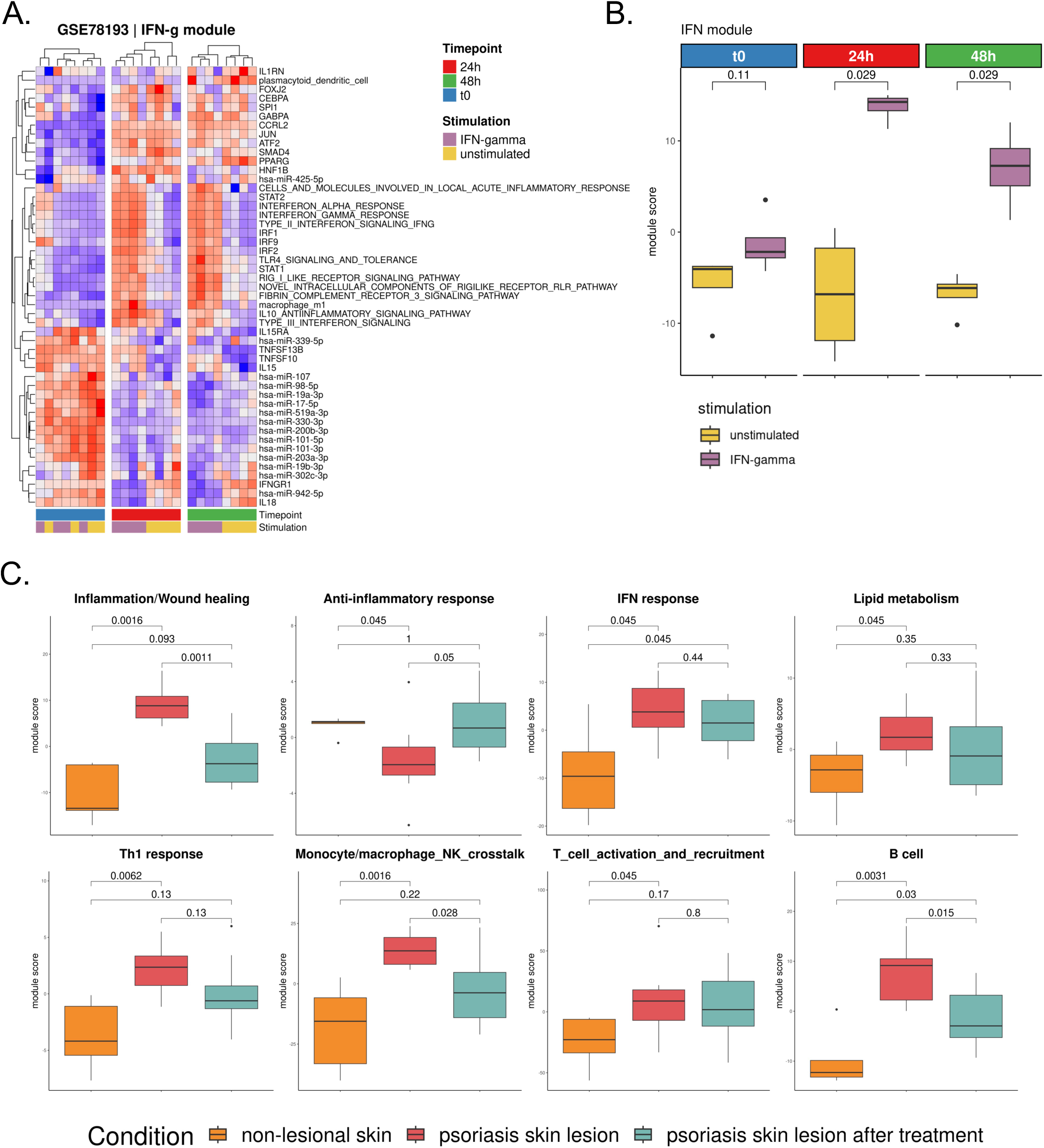
Modules Capture Immune and Inflammation-Related Biological Mechanisms. **(A)** Validation of module interpretation on a public perturbation dataset. Heatmap shows the activity of IFN module features across healthy blood samples, either treated with IFN-γ or left untreated, followed across three time points. Red indicates upregulated activity, blue indicates downregulated activity. Yellow represents IFN-treated samples, and purple represents unstimulated samples. **(B)** Boxplots show differences in module activity between stimulated and unstimulated samples at the three time points. **(C)** Comparison of module activities in the GSE249936 dataset, comparing healthy skin, psoriatic skin, and psoriatic skin after dithranol treatment. Wilcoxon-test p-values were computed, and only modules with at least one significant comparison (p<0.05) are shown.

Upon IFN-γ stimulation, immune modules displayed distinct patterns of responsiveness (**Supplementary Figure 10**). As expected, the IFN module showed the strongest and most consistent modulation across both 24h and 48h, confirming its specificity as a readout of IFN-driven perturbation. Most other modules remained largely unchanged, including B_cell, Lipid_metabolism, T_cell_activation_and_recruitment, and Lymphoid_cell_development, while processes such as ECM remodeling and Impaired cell proliferation were unaffected. A notable exception was the TLR/NFκB_response/innate_immune_response module, which exhibited a modest but significant increase at 48h (Wilcoxon-test, p=0.029), consistent with downstream cross-talk between IFN signaling and innate immune activation. Strikingly, the Monocyte/macrophage_NK_cell_crosstalk module also responded strongly. This module displayed changes nearly as pronounced as those of the IFN module, indicating that IFN-γ stimulation engages not only direct IFN-driven pathways but also innate immune circuits centered on monocyte - NK interactions. This observation was consistent with experimental evidence that coculture of monocytes or macrophages with NK cells induces NK cell activation, characterized by upregulation of CD69, enhanced IFN-γ secretion, and cytotoxic degranulation, particularly when monocytes/macrophages are exposed to microbial or inflammatory stimuli.^63^ Thus, the responsiveness of the Monocyte/macrophage_NK_cell_crosstalk module in our perturbation setting highlighted the ability of our framework to capture both primary IFN signaling and the downstream activation of innate cell–cell communication networks.

These results confirmed that our modules capture coherent immune responses, supporting their use to monitor pathway activation and immune dynamics.

### Immune Modules Reveal Shared and Unique Processes in Autoimmune Diseases

Our modules grouped co-regulated immune features into interpretable biological programs to capture immune dysregulation across autoimmune diseases. To quantify the enrichment of each module across all autoimmune diseases, we calculated the area under the ROC curve (AUC) for each module comparing disease affected to healthy samples across all datasets. We performed this analysis across all blood- and tissue-derived samples, which represent our discovery and validation cohorts respectively. We assessed module enrichment as the average AUC score per disease across all studies for each disease. We also evaluated consistency by computing the inverse of the standard deviation of AUCs, providing a stability-weighted assessment of each module discriminative power (**Figure 4B**). Our analysis revealed both broadly conserved and disease-specific immune perturbations.

The IFN response module emerged as a strong pan-autoimmune biomarker for multiple diseases (mean_AUCs_blood_=0.6 ; mean_AUCs_tissue_=0.84) with strong and consistent enrichment in SLE across both blood (mean_AUC=0.79, integrated p<2.2e-16) and skin (mean_AUC=0.98, integrated p<2.2e-16), reflecting the central role of type I IFNs in SLE pathogenesis.^17,19,20,35,44,61^ Other immune processes like inflammation/wound healing, TLR/NF-κB signaling, and endothelial dysfunction were also significantly enriched across multiple diseases, such as SSc (e.g., SSc skin TLR/NFκB_response/innate_immune_response module mean_AUC=0.70, p<0.05).^64^

Interestingly, modules not traditionally associated with immune function also emerged as relevant in disease-specific contexts. For example, we observed significant enrichment of the Lipid_metabolism module in T1D (mean_AUC=0.66, p<0.05), consistent with the metabolic dysregulation in T1D.^65,66^

In contrast, several modules were selectively downregulated in disease. Most notably, the ECM remodeling module was strongly repressed in salivary glands of SS patients (mean_AUC=0.18, p<0.05), suggesting a lack of active matrix turnover or tissue repair despite the presence of inflammation. Although fibrotic changes have been described in SS,^67^ particularly in advanced or atrophic glands, these may reflect localized or late-stage events not consistently captured in bulk transcriptomic profiles across patients. The absence of robust ECM remodeling signatures in most samples may therefore indicate that fibrosis in SS may develop through mechanisms that may be temporally or spatially restricted, or may involve post-transcriptional regulation not detectable in our analysis. In SSc, by contrast, ECM remodeling was a recognized pathological hallmark.^68,69^ Our analysis shows a significant but modest enrichment in SSc skin (mean_AUC=0.56, p<0.05), indicating that while matrix-related genes were more active during disease, this process was not consistent across studies.

Our analysis provided an overall picture of which biological processes are most relevant in each disease, highlighting both commonalities and differences in their underlying mechanisms.

### Immune Modules Are Broadly Applicable in Capturing Key Autoimmune Processes

To further validate the utility and biological relevance of our modules, we analyzed an independent validation cohort (GSE249936). This dataset profiles skin samples of psoriatic patients, including non-lesional skin, psoriatic lesions, and psoriatic lesions treated with dithranol, a well-established anti-psoriatic agent known for its anti-inflammatory effects.

Several modules showed significant differences between conditions (Wilcoxon-test, p<0.05) (**Figure 5C**). The Inflammation/Wound_healing module, in particular, exhibited significant downregulation in psoriatic lesions (Wilcoxon-test, p=0.0011) after dithranol treatment, consistent with its known anti-inflammatory properties.^70,71^

Beyond inflammation, we observed elevated activity of the IFN response module in psoriatic lesions compared to non-lesional skin (Wilcoxon-test, p=0.045, **Figure 5C**), in agreement with known activation of IFN pathways in Ps.^72^ Concomitantly, we observed significant changes in the Monocyte/macrophage_NK_cell_crosstalk module (Wilcoxon-test, p<0.05, **Figure 5C**), reflecting changes in cellular interactions critical to the inflammatory environment of psoriatic skin.^73^

### Immune Modules Predict Treatment Response in Crohn’s Disease and Ulcerative Colitis

After validating the ability of our immune modules to capture underlying biological changes, we next assessed their clinical utility to predict patient outcomes.

We analyzed transcriptomes (GSE16879) of gut mucosal biopsies from 61 IBD patients (24 with UC, 19 with Crohn’s colitis (CDc), and 18 with Crohn’s ileitis (CDi)) and 12 control patients with normal mucosa (6 from the colon and 6 from the ileum). These patients, refractory to corticosteroids and/or immunosuppression, underwent endoscopic biopsies of actively inflamed mucosa before and 4-6 weeks after their first infliximab infusion. Infliximab is a monoclonal antibody that targets tumor necrosis factor-α (TNF-α), thus reducing inflammation and promoting mucosal healing.^74^ Patients were classified as responders or non-responders to infliximab based on endoscopic and histologic findings at 4-6 weeks post-treatment.

Starting from post-treatment samples, we quantified the Inflammation/Wound_healing module activity, as infliximab is known to target inflammatory pathways central to IBDs.^74,75^ We observed a significant (Wilcoxon-test, p=6.3e-05 for CD ; p=2.2e-03 for UC) (**Figure 6A**) reduction in module activity in responders compared to non-responders, consistent with the suppression of inflammatory processes and restoration of tissue repair mechanisms following successful treatment.

**Figure 6.**
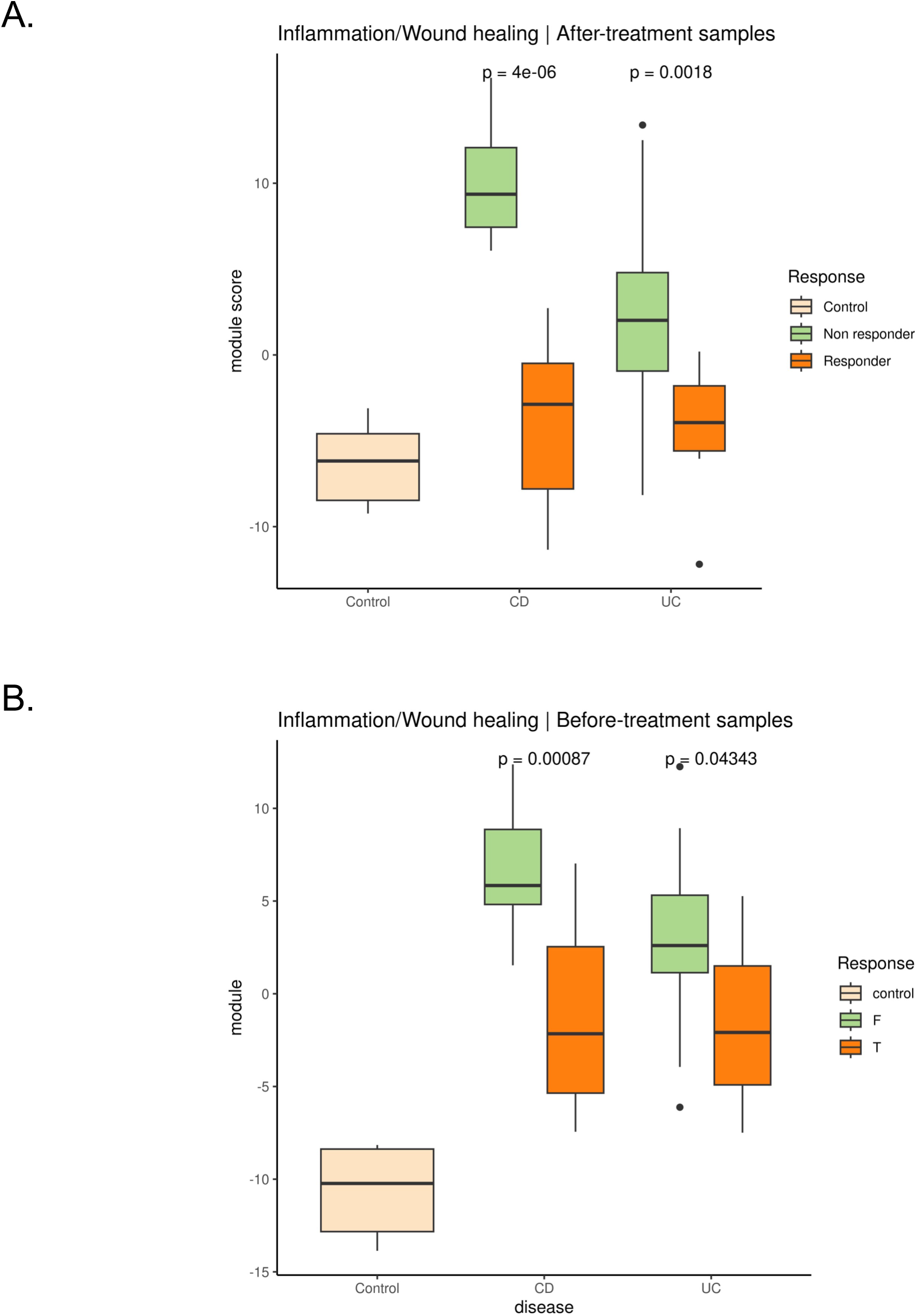
Inflammation Module Activity Distinguishes Treatment Response in IBD. **(A)** In colonic tissue biopsies collected after infliximab treatment, the inflammation/wound healing module score was significantly lower in responders (orange) compared to non-responders (green) in both Crohn’s disease (CD) and ulcerative colitis (UC). **(B)** In baseline (pre-treatment) samples from the same cohort, future responders (orange) already showed lower module scores compared to non-responders (green), indicating the module’s potential to predict therapeutic response. Wilcoxon-test p-values are shown. Control samples (beige) are included as reference.

Next, we determined whether any of our modules could distinguish responders from non-responders before treatment. Consistently with infliximab mechanism of action, we observed significantly reduced activity of the inflammatory module in responders compared to non-responders at baseline in both CD (Wilcoxon-test, p=0.0037) and UC (Wilcoxon-test, p=0.0337) patients (**Figure 6B**). This finding suggests that baseline inflammatory activity, as captured by the module, may serve as a predictive marker for therapeutic response.

To further validate the potential of our modules to predict infliximab response at baseline, we analyzed two additional independent cohorts profiling IBD patients before and after treatment. The first cohort (GSE12251) includes 22 IBD patients who underwent colonoscopy with biopsies collected before starting infliximab treatment. The second cohort (GSE14580) comprises 24 patients with active UC, also treated with infliximab, and 6 healthy controls. Baseline module activity scores accurately predicted responder status at baseline in both cohorts (GSE14580, AUC=0.742 ; GSE12251, AUC=0.803, **Supplementary Figure 11**).

Overall, our results demonstrate the robustness of the inflammation module in capturing pre-treatment immune activity associated with therapeutic outcomes, highlighting its potential as a predictive biomarker for treatment response in UC.

### Module Activity Correlates with Disease Severity in Psoriasis

Finally, we assessed the ability of our modules to predict disease severity in autoimmune disorders.

We analyzed a second Ps cohort (GSE117239) of transcriptomes from patients treated with either ustekinumab, an IL-12 and IL-23 inhibitor, or etanercept, a TNF inhibitor. This cohort included samples from lesional and non-lesional skin biopsies collected at baseline and after 12 weeks of treatment, paired with Psoriasis Area and Severity Index (PASI) scores and response status. Higher PASI scores indicate more severe disease conditions, correlating with the extent and severity of psoriatic lesions.

At baseline, both responders and non-responders exhibited similar PASI scores across treatment groups (Wilcoxon-test, p=0.17, p=0.18, p=0.16 for etanercept, ustekinumab 45 mg and ustekinumab 90 mg, respectively), reflecting comparable disease severity before therapy. However, after 12 weeks of treatment, significant reductions in PASI scores were observed in responders treated with ustekinumab (both 45 mg and 90 mg) (Wilcoxon-test, p=1.6e-05 and p=1.9e-08, respectively) and etanercept (Wilcoxon-test, p=1.5e-10). In contrast, non-responders showed lower reduction in PASI scores, indicating lack of therapeutic benefit (**Figure 7A**).

**Figure 7.**
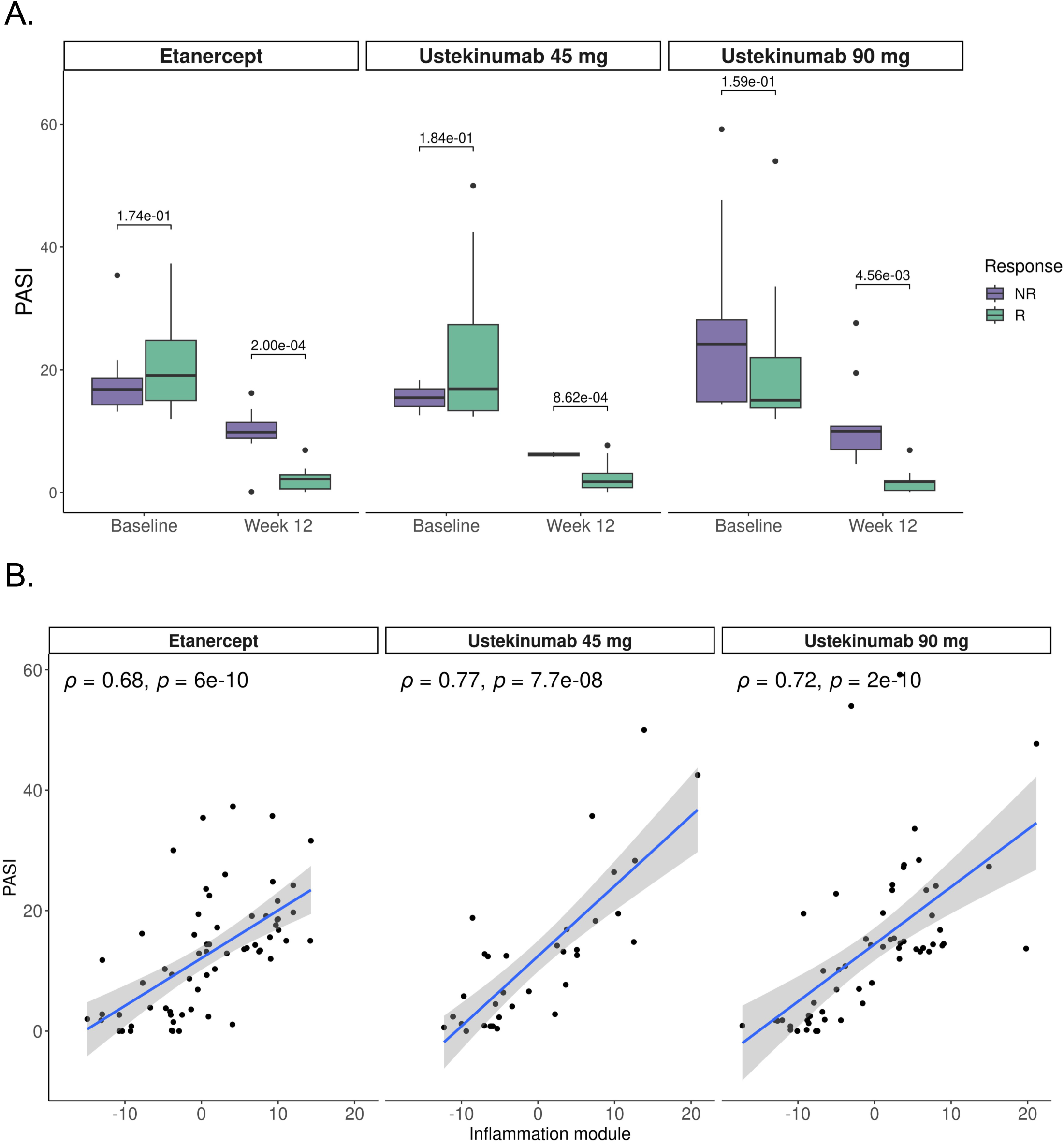
Module Activity Correlates with Clinically Determined Disease Severity Score in Psoriasis. **(A)** Boxplot of PASI scores for patients at baseline and after 12 weeks of treatment with Etanercept, Ustekinumab 45 mg, and Ustekinumab 90 mg. P-values were calculated by Wilcoxon-test. **(B)** Correlation plots showing the relationship between inflammation module activity and PASI scores measured by Spearman’s correlation and its corresponding significance.

Because ustekinumab and etanercept are anti-inflammatory drugs,^76^ we investigated whether the Inflammation/Wound_healing module was associated with disease severity by computing module activity scores for all samples in the dataset and correlating them with PASI scores. Remarkably, we observed strong positive correlations between the inflammation module scores and PASI scores across etanercept-(R=0.6, p=6e-10) and ustekinumab-(R=0.77, p=7.7e-8, 45 mg dosage; R=0.72, p=2e-10, 90 mg dosage) treated patients (**Figure 7B**), demonstrating that inflammation module heightened activity was closely associated with more severe disease. This underscores the clinical relevance of the inflammation module, not only as a marker of inflammatory processes but also as a robust correlate of disease activity, further validating the clinical utility of our modules.

Taken together, these findings provide an integrative immunogenomic framework that unifies diverse datasets across tissues and diseases, delineates common and disease-specific immune modules, and highlights key processes such as IFN signaling and inflammation, thereby offering a comprehensive view of the immune landscape in autoimmunity with clear biological and clinical relevance.

## Discussion

We conducted a comprehensive immunogenomic analysis of 10 autoimmune diseases by integrating 13,263 public transcriptomes from blood and tissue from patients and healthy controls. Our goal was to develop a cross-disease framework to identify and study shared and disease-specific immunological mechanisms across autoimmunity.

Our analysis of blood transcriptomes revealed that diseases with distinct clinical and histopathological manifestations can nevertheless converge at the molecular level. For example, CD and UC are unique entities within the IBDs spectrum, with differences in anatomical involvement, histological depth of inflammation, and endoscopic appearance^77^. Despite this heterogeneity, we observed that their blood expression profiles are remarkably similar, pointing to a shared transcriptional program underlying intestinal autoimmunity. Our result mirrors genetic studies showing substantial overlap between the two conditions^16^, and suggests that systemic immune responses in CD and UC may be more similar than their clinicopathological definitions would imply.

A similar trend was observed across systemic autoimmune rheumatic diseases (SLE, RA, SS, and SSc). Despite clinical diversity, they are characterized by the presence of autoantibodies, profound immune system dysregulation, and shared genetic susceptibility across the HLA region.^17^ Concordantly, our transcriptional analysis revealed that these four diseases cluster tightly together in blood, despite their clinical presentations. Our analysis of circulating transcriptional programs is suggestive of common pathogenic mechanisms across systemic autoimmune rheumatic diseases.

Conversely, the transcriptional profile of JIA was remarkably different from other conditions, suggesting the presence of disease-specific molecular mechanisms and hinting that the etiology of JIA may be more strongly shaped by unique genetic determinants. Indeed, numerous studies have investigated the genetic basis of JIA and identified a range of risk loci^78,79^.

Globally in blood, we observed a consistent upregulation of IFN and antiviral response pathways, alongside a broader pattern of innate immune activation coupled with impaired adaptive modulation. These findings suggest a shared molecular architecture across autoimmune diseases, with the potential of pan-disease therapeutic avenues.

Since peripheral blood does not provide a direct view on tissue immunology, we included 7,025 tissue transcriptomes in our analysis. Overall, we identified both shared and disease-specific autoimmune transcriptional programs across tissues. Diseases affecting the same compartment, such as CD and UC, showed similar expression profiles, while systemic rheumatic diseases displayed more distinct transcriptional signatures. SLE provided a clear example of context-dependence: LN displayed a transcriptional profile distinct from cutaneous lupus, underscoring that even within the same disease, molecular programs diverge according to the affected organ.^80^

Our analysis confirmed a general enrichment of IFN and inflammatory pathways, and uncovered context-specific alterations, such as downregulation of genes involved in ECM remodeling and tissue integrity,^41–43^ and complement activation in LN^45^. These tissue and disease-specific pathways suggest that the pathobiology of autoimmunity impair local tissue structure and function beyond immune infiltration.

To better characterize the role of the immune system in autoimmunity, we extracted biologically meaningful immune features from transcriptomics data. Our higher-order features capture distinct layers of immune regulation, providing a systems-level perspective that both complemented and extended conventional differential expression analyses. Features-based clustering revealed finer granularity than gene-level patterns, exposing previously unappreciated distinctions among diseases and refining disease relationships. In particular, we observed strong similarities between SLE and RA - an association well established by genetic and transcriptomic studies.^37,38^ Our approach also recapitulated known immunopathological hallmarks, including lymphopenia in SLE,^32,33^ M1 macrophage enrichment across inflammatory diseases,^34^ and pervasive type I IFN activity across autoimmunity.^44,61,81^ Together, these findings demonstrate that immune feature abstraction yields a more integrated and mechanistically interpretable view of autoimmune biology, while capturing cross-disease relationships with greater resolution than gene-level analyses alone.

Our work provides, to our knowledge, the first systematic comparison of peripheral and tissue-level immunity across multiple autoimmune diseases. Blood and tissue compartments revealed only limited concordance, with partial overlap across both differentially regulated genes and features. These differences were disease-specific: CD and UC displayed high concordance between blood and gut-derived transcriptomes, whereas RA and SS showed marked divergence. This confirms that while peripheral blood reflects broad systemic immune activation, it may fail to fully capture the local transcriptional programs driving pathology within target organs. Nevertheless, several immune features - notably M1 macrophages, pDCs, and IFN pathway activity - were consistently enriched across both compartments. In particular, we detected strong co-regulation between type I IFN modules and pDCs abundance, consistent with the role of pDCs as the major source of IFN and their established contribution to SLE pathogenesis.^60,62^ Our results indicate that this IFN–pDC axis extends beyond SLE and represents a recurring inflammatory circuit across autoimmune diseases. Together, these findings highlight immune features that bridge systemic and tissue-level contexts and underscore the central role of IFN-driven innate activation as a unifying mechanism in autoimmunity.

We organized our immune features into 15 immune modules to compare immune processes at a higher resolution. These modules captured molecular and cellular processes reflecting either direct immune responses - such as IFN signaling, inflammatory activation, and lymphocyte regulation - or a broader disease milieu - such as fibrosis, tissue remodeling, and metabolic reprogramming.

We first successfully validated our IFN module in whole blood using transcriptional data from IFN stimulation experiments. We then measured module activity across diseases and compartments, uncovering coherent and biologically interpretable results. We found the IFN module to be broadly engaged across conditions, with particularly strong signals in SLE, SS, and SSc - in line with the pervasive interferonopathy reported in these diseases.^35,57,61^ In contrast, the T-cell activation/recruitment module showed modest contribution in SLE blood samples, consistent with the well-described lymphopenia in SLE,^32,33^ while it was markedly enriched in SLE tissues-derived transcriptomes, suggesting active recruitment of T cells from the circulation into inflamed organs. B-cell–related activity also increased at disease sites, echoing the central role of local antibody production in autoimmunity.^82,83^ Tissue effects were also evident for modules reflecting inflammation/wound healing and innate immune activation, which dominated in affected organs and are consistent with previous observations of tissue-destructive inflammatory cascades.^84^ Similarly, the integrin/adhesion module was strongly represented in tissue but not in blood, aligning with the established role of adhesion and extravasation pathways in immune cell trafficking.^85^

Together, these results extend our knowledge of the differences between systemic and local immune programs: systemic processes, such as IFN activation, are readily captured in blood, while tissue-specific processes - including adhesion, innate activation, and tissue remodeling - become prominent only at the site of disease. Our framework helps explain why blood transcriptional profiling reflects broad immune activation yet misses the full extent of local pathology, providing new potential circulating biomarkers and tissue-targeted therapeutic targets.

The clinical and predictive potential of our immune modules extends their utility beyond descriptive biology. By integrating diverse immunological signals into coherent units, the modules capture dimensions of immune activity that are directly relevant to patient outcomes. In validation, modules consistently reflected biologically meaningful changes following treatment. For example, the Inflammation/Wound_healing module decreased in activity after exposure to anti-inflammatory therapies, mirroring suppression of tissue inflammation and restoration of repair processes. While prior studies have shown that inflammatory signatures decline with effective therapy in IBDs and Ps,^86,87^ our framework provides a more integrated and interpretable readout, capturing these shifts as coordinated immune programs rather than as isolated gene-level changes.

Beyond measuring treatment effects, our results demonstrate that immune modules carry a predictive role. In IBDs, baseline activity of the Inflammation/Wound_healing module distinguished infliximab responders from non-responders reproducibly across multiple independent cohorts.

Similarly, in Ps we observed that the inflammation module activity correlated strongly with the Psoriasis Area and Severity Index (PASI), a gold-standard clinical parameter of disease burden. This result directly links molecular activity to clinical severity, emphasizing the value of our framework for bridging transcriptomic and clinical domains. While previous transcriptomic studies in Ps have reported associations between PASI and selected inflammatory pathways,^88^ our approach improves upon previous approaches by capturing inflammation as a module-level process, increasing reproducibility and interpretability across patients and cohorts. Moreover, the ability of modules to track therapeutic changes further supports their potential as dynamic biomarkers of treatment response.

Taken together, these findings position immune modules as a powerful framework for clinical translation, providing predictive potential and direct correlation with clinical endpoints. Unlike single-gene biomarkers, which often lack reproducibility, modules across features derived by integrating multiple independent cohorts offer greater reproducibility and accuracy for downstream applications. Our immune modules are both mechanistically relevant and clinically actionable, with the potential to inform patient stratification, guide therapeutic targeting, and monitor treatment efficacy, advancing the path toward precision medicine in autoimmunity.

While this study offers extensive validation and broad disease coverage, it also has limitations. Computational inference of immune features depends on predefined gene sets and models, which may not comprehensively capture the complexity of the underlying biology. Furthermore, while we validated the prediction of our modules in independent datasets, experimental validation will be needed to establish causal relationships and guide therapeutic development. Additional caveats should be noted. First, our analysis relies exclusively on transcriptomic data; integrating multi-omics layers such as proteomics, epigenomics, or single-cell data would provide a richer view of immune regulation and help bridge the gap between transcriptional activity and functional output. Second, although we applied rigorous normalization and meta-analytic approaches, batch effects and differences in study design remain potential sources of confounding. Third, tissue coverage was uneven across diseases, with certain autoimmune conditions represented primarily in blood, which may limit the generalizability of tissue-level findings. Relatedly, most datasets captured static snapshots of disease rather than longitudinal dynamics, constraining our ability to fully model temporal immune trajectories. Fourth, our framework does not directly incorporate clinical covariates such as treatment history, comorbidities, or demographic factors, which may influence immune profiles and merit integration in future analyses. Specifically, although meta-analysis improves generalizability and helps mitigate sample-specific sex biases, gender information was not consistently available across datasets, preventing systematic sex- and gender-based analyses; incorporating this dimension in future studies could provide additional insight into immune regulation and disease mechanisms.

Nonetheless, the interpretability, generalizability, and predictive power of our immune modules underscore their translational potential. Future work integrating multi-omics, prospective longitudinal cohorts, and mechanistic experiments will further refine this framework, advancing its application to biomarker and drug discovery in autoimmunity.

Finally, to support transparency and foster collaboration, we developed an interactive Shiny app that allows users to dynamically explore and download processed data and results. In parallel, we released all code and data through a public GitHub repository. By publicly releasing both the processed datasets and the analytical framework, we provide a valuable resource for the research community - one that others can build upon to further advance our understanding of autoimmunity.

## Methods

### Data collection

In this study, we aimed to compile all available bulk gene expression case-control studies for 10 autoimmune diseases from the Gene Expression Omnibus (GEO) database. Using the utilities provided by the Entrez Direct API, we performed keyword-based searches to identify relevant studies in GEO and retrieved their unique identifiers. Subsequently, we utilized the GEOquery R package to extract metadata associated with each sample from the identified studies. The dataset collection process was finalized in October 2022.

For this analysis, we included studies that met the following criteria: (1) contained both case and control individuals, (2) utilized bulk sequencing technologies (including microarray and RNA-seq), (3) had a minimum of four patients per study, (4) featured samples derived from either whole blood or affected tissue, (5) retained only baseline samples in longitudinal studies to ensure one sample per patient, and (6) excluded samples from patients undergoing treatment, when such information was available. For RNA-seq studies, we included only those that provided TPM-normalized data or raw counts. Table 1 summarizes the number of studies collected for each disease.

A notable challenge with GEO data is the lack of standardized metadata annotation, which limits its direct usability for computational analyses. To address this issue, we undertook extensive manual curation and standardization of metadata across all studies. Specifically, we manually annotated fields such as “series_id”, “geo_accession”, “disease”, “gender”, “age”, “timepoint”, “sample_type”, and “sample_source”, among others, ensuring consistency wherever possible. However, some fields were unavailable for certain studies.

For microarray data, we leveraged the built-in tools of the MetaIntegrator R package used for meta-analysis, which automatically downloads and normalizes microarray datasets. In contrast, for RNA-seq studies, we either used TPM-normalized data directly or performed TPM normalization on raw count data. All datasets were converted to a log scale prior to conducting the meta-analysis.

This effort enabled the creation of a curated dataset of autoimmune studies, harmonizing metadata across a vast number of samples from the GEO database, thereby providing a valuable resource for advancing research in the field of autoimmunity.

The data are available at our GitHub repository (https://github.com/rominaappierdo/AutoimmuneLandscape).

Both data and metadata can be interactively explored through our Shiny app (http://160.80.36.177:4242/myapp_Autoimmune/shinyApp_Autoimmune/).

### Meta-analysis

Meta-analysis is a powerful statistical technique that synthesizes findings from multiple independent studies addressing a common research question, providing a unified and statistically robust framework for interpretation.^15^ Central to this technique is the concept of effect size, a standardized metric that quantifies the magnitude and direction of a relationship, enabling comparisons across studies with varying methodologies and measurement scales. However, the effect size observed in any individual study is only an estimate of the true effect size and may differ from it due to sampling error.

This can be denoted mathematically as:

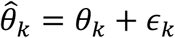

where 𝜃̂_*k*_ represents the observed effect (estimated of the true effect size), 𝜃_*k*_ the true effect of a study 𝑘, and 𝜖_*k*_ the sampling error. This error arises because studies are based on samples rather than entire populations, and its magnitude is inversely related to sample size.

Meta-analysis takes into consideration the precision of effect size estimates. When combining results from multiple studies, they assign higher weight to those with higher precision (i.e., lower sampling error), as these provide a more accurate representation of the true effect.

A typical way to express 𝜖 is by using the standard error (SE). The standard error (SE), calculated as the standard deviation divided by the square root of the sample size, reflects the variability in these estimates, with larger samples producing smaller SE values and thus more precise estimates:

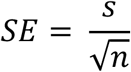

There are different ways in which effect sizes can be computed. Usually, if the goal of the analysis is to compare two groups, the standardized mean difference (SMD) expresses group differences in units of standard deviation, making it particularly useful in studies where outcomes are measured on different scales.

The standardized mean difference is also often called *Cohen’s d* and computed as:

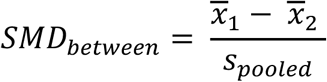

where *S*_pooled_ is:

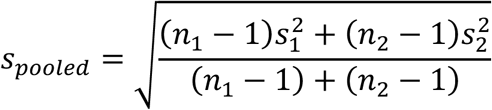

Of note, standardized mean differences are often used in meta-analysis because they can be compared between studies. The standard error of the SMD between two conditions (SMD_between_) can be calculated with:

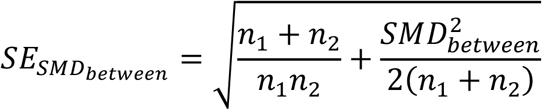

SMDs tend to exhibit an upward bias in studies with small sample sizes. To address this, they are often adjusted, resulting in an effect size known as Hedges’ *g*, which we employed in our study:

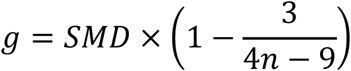

After computing effect sizes for each study, meta-analysis combines them into a single summary effect. To achieve this, a specific statistical model must be assumed. In this project, we employed a random-effects model to account for variability between studies. Differences in study populations, methodologies, or experimental conditions often introduce variability that cannot be explained by sampling error alone. Unlike a fixed-effects model, which assumes that all studies estimate the same true effect and that differences arise solely from random sampling error, a random-effects model acknowledges that studies may differ due to a combination of factors. This model introduces an additional variance component to reflect between-study heterogeneity. Instead of assuming all studies stem from a single homogeneous population, the random-effects model treats each study as an independent sample from a broader “universe” of populations. This framework is particularly appropriate for meta-analyses where variability between studies is expected.

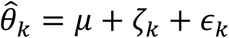

where 𝜃̂_*k*_ is the observed effect size in study 𝑘, 𝜇 is the overall mean effect size, 𝜁 is the random effect representing the deviation of study 𝑘’s true effect from the overall mean due to between-study heterogeneity and 𝜖_*k*_ is the within-study error (sampling error).

This between-study variability, known as heterogeneity, is incorporated into the analysis through an additional variance component (𝜏^*k*^), that represents the variance of the distribution of true effect sizes and adjusts the weighting of each study.

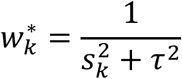

In this study we used the DerSimonian and Laird (DL) method for estimating tau^2^.

By combining within-study variance (standard error) and between-study variance, the random-effects model ensures that studies with greater variability contribute less to the pooled estimate.

The final formula to calculate pooled effect size with random effect model is:

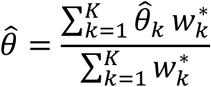

This approach provides a robust representation of real-world data, accommodating the natural differences that exist between studies while enabling broader generalizations.

For the meta-analysis of gene expression datasets, we employed the MetaIntegrator framework (ref), an established computational method specifically designed to integrate and analyze transcriptomic data from multiple independent studies. Meta-analysis of gene expression data is crucial in consolidating findings across different datasets, enabling the identification of robust and reproducible gene signatures associated with the condition of interest, even in the presence of biological and technical variability. MetaIntegrator addresses these challenges by first performing dataset harmonization and quality control, ensuring that all datasets meet baseline standards for inclusion.

Following dataset harmonization, MetaIntegrator calculates effect sizes for each gene across all datasets, using standardized mean differences with correction for small sample bias (Hedge’s g) to quantify the magnitude and direction of association between gene expression and disease state. To address inter-study heterogeneity the method uses a random-effects model.

Gene signatures are then identified based on combined effect sizes and significance thresholds, prioritizing genes with consistent associations across studies. To enhance biological interpretability, the MetaIntegrator method includes tools for functional annotation and pathway enrichment analysis, enabling further insights into the biological processes and pathways represented by the identified gene signatures. This rigorous approach provides a robust and reproducible set of candidate genes for further validation and exploration, particularly valuable for complex diseases where single-cohort studies may lack sufficient power.

### Over-representation analysis

To investigate the biological processes associated with differentially regulated genes, we performed overrepresentation analysis (ORA) using the clusterProfiler R package.^89^ ORA identifies pathways or functional categories that are significantly enriched within a given gene set compared to a background set of all expressed genes. For this study, we focused on the Gene Ontology (GO) Biological Processes database, which provides a structured and comprehensive view of biological functions and processes. By inputting the lists of upregulated and downregulated genes for each condition, the algorithm compares the observed number of genes in each pathway to the expected number based on the background set, calculating enrichment scores and statistical significance using methods such as Fisher’s exact test or hypergeometric distribution. The results highlight pathways that are disproportionately represented in the gene sets, providing a deeper understanding of the molecular mechanisms underlying autoimmune diseases.

### Transcriptomic Features

#### Cell mixture deconvolution

Cell-mixture deconvolution is a computational technique used to estimate the relative proportions of distinct cell types within a bulk tissue sample. This method dissects complex biological samples into their cellular components, offering a detailed view of cellular heterogeneity that would otherwise remain obscured in bulk RNA-sequencing data. In this study, we applied the immunoStates basis matrix, a reference framework comprising gene expression profiles from well-characterized immune cell types, including T cells, B cells, monocytes, and natural killer cells.^90^

The deconvolution process leverages marker genes selectively expressed by specific cell types. By comparing the expression levels of these markers in the bulk sample to those in the reference matrix, a linear modeling framework estimates the proportions of each cell type. This approach transforms averaged gene expression data from bulk RNA-sequencing into interpretable contributions from individual immune cell types.

By focusing on immune cells using the immunoStates matrix, this analysis enabled us to quantify shifts in cellular composition linked to pathological states. This methodology is integral to understanding the cellular mechanisms driving disease progression and offers insights that may inform the development of targeted therapeutic strategies.

#### ssGSEA

Single-sample Gene Set Enrichment Analysis (ssGSEA) is an algorithm used to compute a single activity score for a predefined gene set within a single sample,^91,92^ offering an interpretable measure of biological processes. Gene sets can represent diverse biological entities, such as pathways (based on genes in the pathway), TFs (via their regulons), or miRNAs (through their target genes). This enables ssGSEA to capture various regulatory and functional activities in a sample-specific manner.

The algorithm operates by ranking all genes in a sample based on their expression levels. For a given gene set, ssGSEA calculates an enrichment score by comparing the distribution of genes in the set to those outside it. Specifically, it uses cumulative distribution functions (CDFs) of the ranked gene expression values: one CDF is computed for genes in the set and another for genes not in the set. The enrichment score is derived as the maximum deviation between these two distributions, quantifying whether the genes in the set are consistently enriched at the top (upregulated) or bottom (downregulated) of the ranked list. This score is normalized to account for differences in gene set size, allowing for fair comparisons across sets. For this study, we applied ssGSEA using the GSVA package in R, which provides a robust implementation of the algorithm. By leveraging ssGSEA, we computed activity scores for pathways, TFs, and miRNA-target sets across our samples.

For this analysis, we curated gene sets from reliable databases, such as pathway gene sets from the Molecular Signature Database (MSigDB) collections (Hallmark, KEGG, and WikiPathways), TF regulons from DoRothEA, and miRNA-target gene sets from miRTarBase.

#### HGNC

To include cytokine, chemokine, and receptor gene expression as a feature, we derived it as a subset from our bulk gene expression dataset, focusing specifically on genes involved in cytokine and chemokine signaling due to their central role in inflammation and immunity. To generate a comprehensive and relevant list of genes, we performed a keyword-based search using the HGNC (HUGO Gene Nomenclature Committee) web server, a well-curated resource for gene nomenclature and classification. By searching for terms such as “cytokines,” “chemokines,” and “interleukins,” we identified 275 genes representing cytokines, chemokines, and their associated receptors. This targeted approach enabled us to systematically analyze the expression and regulation of these pivotal immune mediators across autoimmune diseases, offering valuable insights into their contributions to disease pathology We manually curated a list of 275 cytokines genes from the HGNC database. This subset was specifically chosen to delve deeper into the gene expression patterns of these molecules, given their pivotal roles in inflammatory and immune processes.

### Module Identification and Biological Interpretation

We computed pairwise Spearman’s correlations between all features, identifying groups of features, or *modules*, that exhibited similar behavior across diseases. Features within the same module are likely involved in related biological processes, providing a more structured and interpretable framework for exploring the autoimmune landscape. These modules provide a structured view of the autoimmune landscape, enabling us to link features to broader processes and gain deeper insights into the molecular underpinnings of these diseases.

To organize the 700 significant features into meaningful groups, we cut the dendrogram to obtain 15 distinct clusters of features. The identification and interpretation of these clusters were primarily driven by manual curation. For each module, we examined the features it contained, reviewed relevant literature, and assigned a biological interpretation based on the shared characteristics of the features within the module. This approach ensured that each module represented a coherent and biologically relevant entity, reflecting processes or pathways likely involved in autoimmune disease mechanisms. Therefore, by combining data-driven clustering with thorough literature-based analysis, we derived modules that provide a structured and meaningful perspective on the autoimmune landscape.

## Validation using Independent Perturbation and Clinical Expression Datasets

### Computation of Aggregate Module Scores per Sample

Since a module consists of multiple features, we first computed individual activity, abundance, or expression scores for each feature within a given sample, depending on the feature type. This resulted in a set of values representing the module’s overall behavior in that sample. To derive a single representative score for each module, we aggregated these values by calculating the sum of activity scores across all its features. Prior to aggregation, feature values were scaled across samples in the dataset to ensure comparability. This approach provides a concise yet biologically meaningful summary of module activity in the single sample, while preserving the contributions of individual features.

### Permutation-Based Validation of IFN Module Specificity in Perturbation Data

To confirm the robustness of our findings and ensure that the IFN module reflects a real biological signal, we performed a control analysis using a permutation test. Specifically, we created 1,000 random modules, each of the same size as the IFN module (49 features), by randomly selecting features from the other 14 modules. This was done separately for each time point in the perturbation experiment to account for temporal variation in the data. For each of these random modules, as well as for the real IFN module, we calculated an effect size, which measures the strength of the observed signal. To determine if the IFN module’s effect size was significantly different from those of the random modules, we transformed its effect size into a z-score, comparing it to the distribution of effect sizes from the 1,000 random modules. We then calculated p-values to quantify the likelihood that the observed result for the IFN module could occur by chance.

### Cross-Compartment Assessment of Immune Module Contribution to Disease Classification

To evaluate the contribution of each module to individual autoimmune diseases across blood and tissue, we assessed their ability to distinguish disease samples from controls. For each module and study within our dataset, we generated Receiving Operator Characteristic (ROC) curves using scores calculated for each module. From there, we computed the area under the curve (AUC) for each module. The AUC measures the discriminatory power of a score (in this case, the module activity score) in separating two groups, with values ranging from 0.5 (no discrimination) to 1 (perfect discrimination). This calculation allowed us to quantify the extent to which each module’s activity was associated with disease status across studies.

This analysis was performed independently for both blood and tissue datasets. For each disease, we aggregated the AUC values from all studies by calculating their mean, providing a summary measure of each module’s overall contribution to disease versus control classification. To further assess the consistency of module performance within each disease, we calculated the standard deviation (SD) on the AUC values. This metric allowed us to evaluate the heterogeneity across studies, where a lower SD indicates greater consistency in module activity’s ability to distinguish disease samples from controls. By combining mean AUC values with heterogeneity assessments, this analysis provided a detailed overview of the relevance of each module across diseases. This approach highlighted the modules that are broadly consistent across studies and those with disease-specific variations, offering insights into the immune processes that define each autoimmune condition.

### ROC Curve and AUC Analysis for Treatment Response Prediction

To validate the predictive capability of the Inflammation/Wound_healing module, we generated ROC curves on our independent validation datasets profiling UC patients before treatment, with known responder and non-responder classifications. We used the AUC of each ROC to quantify each module’s ability to distinguish between responders and non-responders based on the module activity scores of individual patients. AUC was calculated as described above.

## Resource availability

### Lead contact

Requests for further information and resources should be directed to and will be fulfilled by the lead contact, Pier Federico Gherardini.

### Materials availability

This study did not generate new unique reagents.

### Data and code availability

- All datasets used in this study are publicly available at https://www.ncbi.nlm.nih.gov/geo/.
- Processed data and the code used for analysis are available on GitHub at https://github.com/rominaappierdo/AutoimmuneLandscape. Additional details and usage instructions are provided in the repository README file.
- A Shiny web application for interactive exploration of the results is accessible at http://160.80.36.177:4242/myapp_Autoimmune/shinyApp_Autoimmune/.
- Any additional information required to reanalyze the data reported in this paper is available from the lead contact upon request.

## Author contributions

Conceptualization: P.F.G., F.V.

Methodology: R.A.

Software: R.A.

Validation: R.A.

Formal Analysis: R.A.

Investigation: R.A.

Resources: R.A.

Data Curation: R.A.

Writing - Original Draft: R.A.

Writing - Review & Editing: R.A., P.F.G., F.V., M.S., G.P., M.H.-C.

Visualization: R.A.

Supervision: P.F.G., F.V., M.S.

Funding Acquisition: M.H.-C.

## Acknowledgements

This study was in part supported by European Union – NextGenerationEU: National Center for Gene Therapy and Drugs based on RNA Technology, CN3 – (code:CN00000041) and NIAMS P30 AR070155.

