## Supplementary_Figures for "The Integrated Cellular and Molecular Landscape of Autoimmunity"

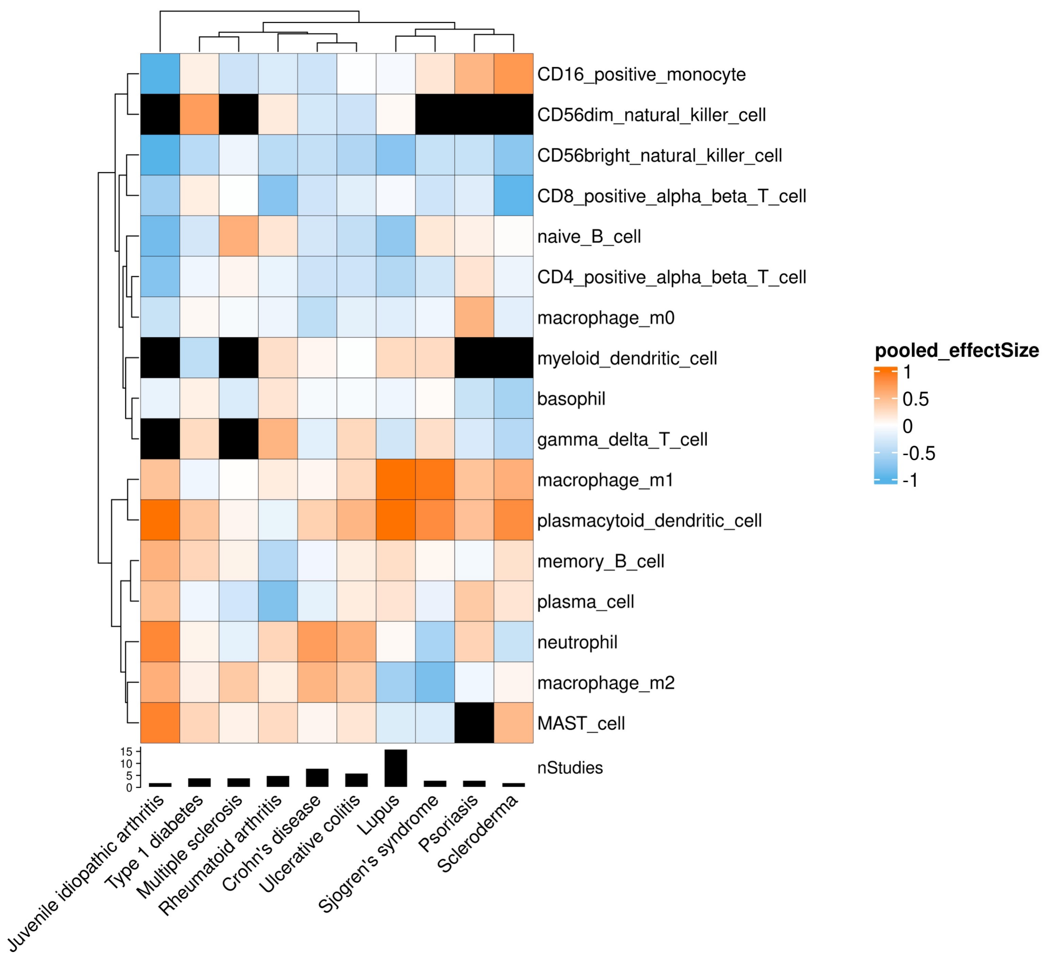


**Supplementary Figure 1.**

Meta-analysis of immune cell proportion changes across autoimmune diseases in blood: each row represents an immune cell type estimated via cell-mixture deconvolution; each column corresponds to one of the ten autoimmune diseases analyzed. The heatmap color indicates pooled effect sizes for the case-control comparisons of estimated cell-type proportions across studies, with positive values (orange) indicating increased abundance in disease and negative values (blue) indicating depletion. Black boxes indicate insufficient data for reliable estimation.

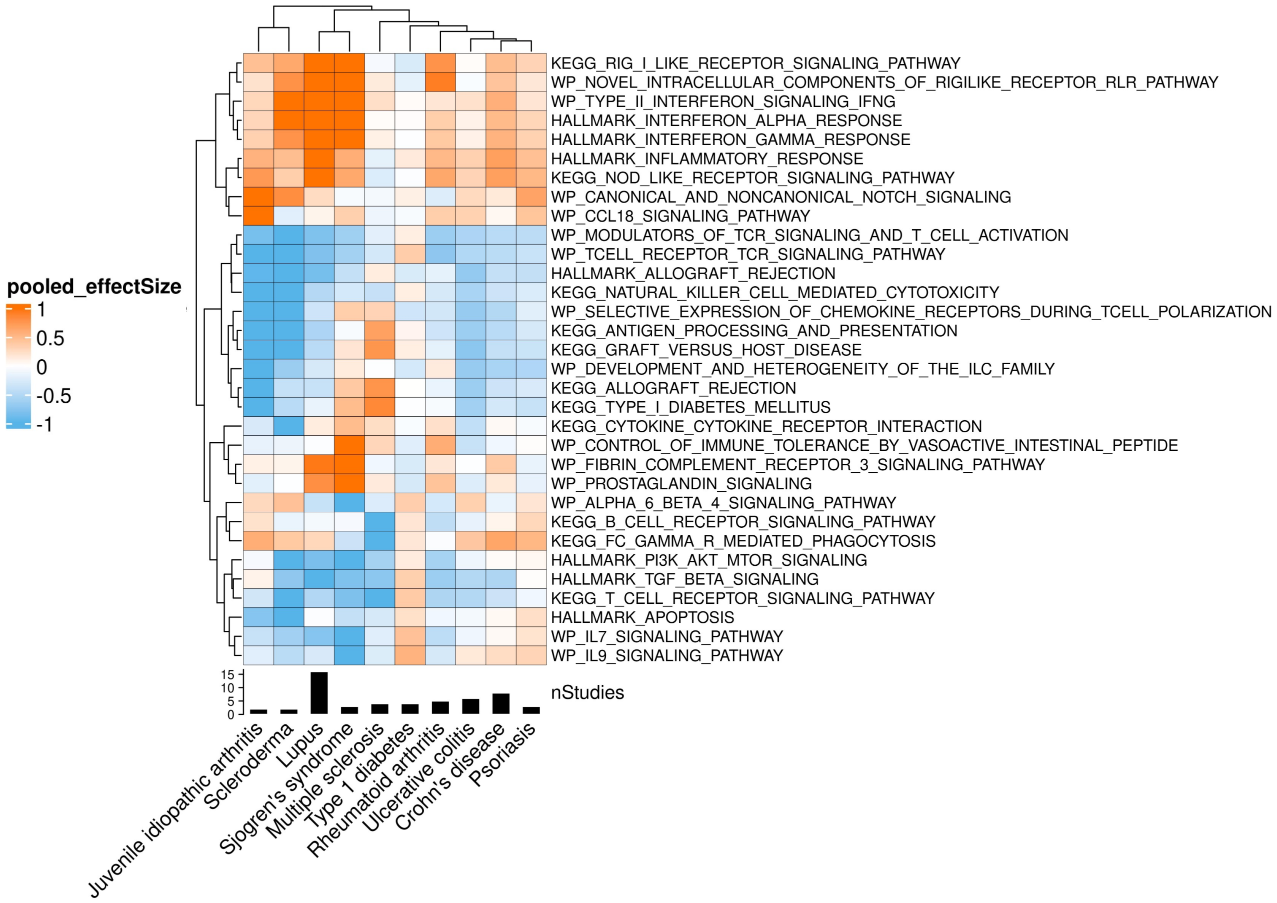


**Supplementary Figure 2.**

Meta-analysis of pathway activity changes across autoimmune diseases in blood: Heatmap displays pooled effect sizes for 30 immune-related pathways estimated via ssGSEA across ten autoimmune diseases. Rows represent pathways derived from MSigDB Hallmark, KEGG, and WikiPathways databases, while columns correspond to diseases. Positive values (orange) indicate increased pathway activity in disease relative to control; negative values (blue) indicate decreased activity. Black tiles mark pathways not assessed due to insufficient data.

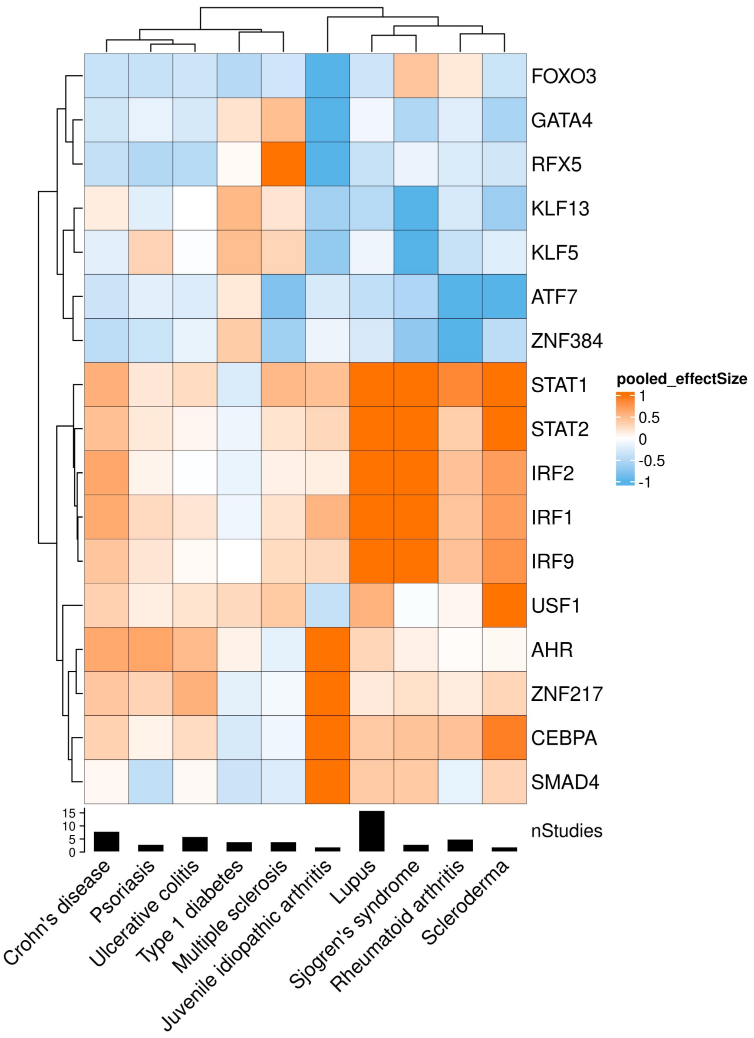


**Supplementary Figure 3.**

Meta-analysis of transcription factor activity across autoimmune diseases in blood: the heatmap shows the pooled effect sizes for 18 transcription factors (TFs) across ten autoimmune diseases, estimated using ssGSEA and DoRothEA regulon-based inference. Rows represent TFs, and columns represent diseases. Positive values (orange) indicate increased activity in disease versus control; negative values (blue) indicate decreased activity. Black tiles indicate insufficient data for analysis.

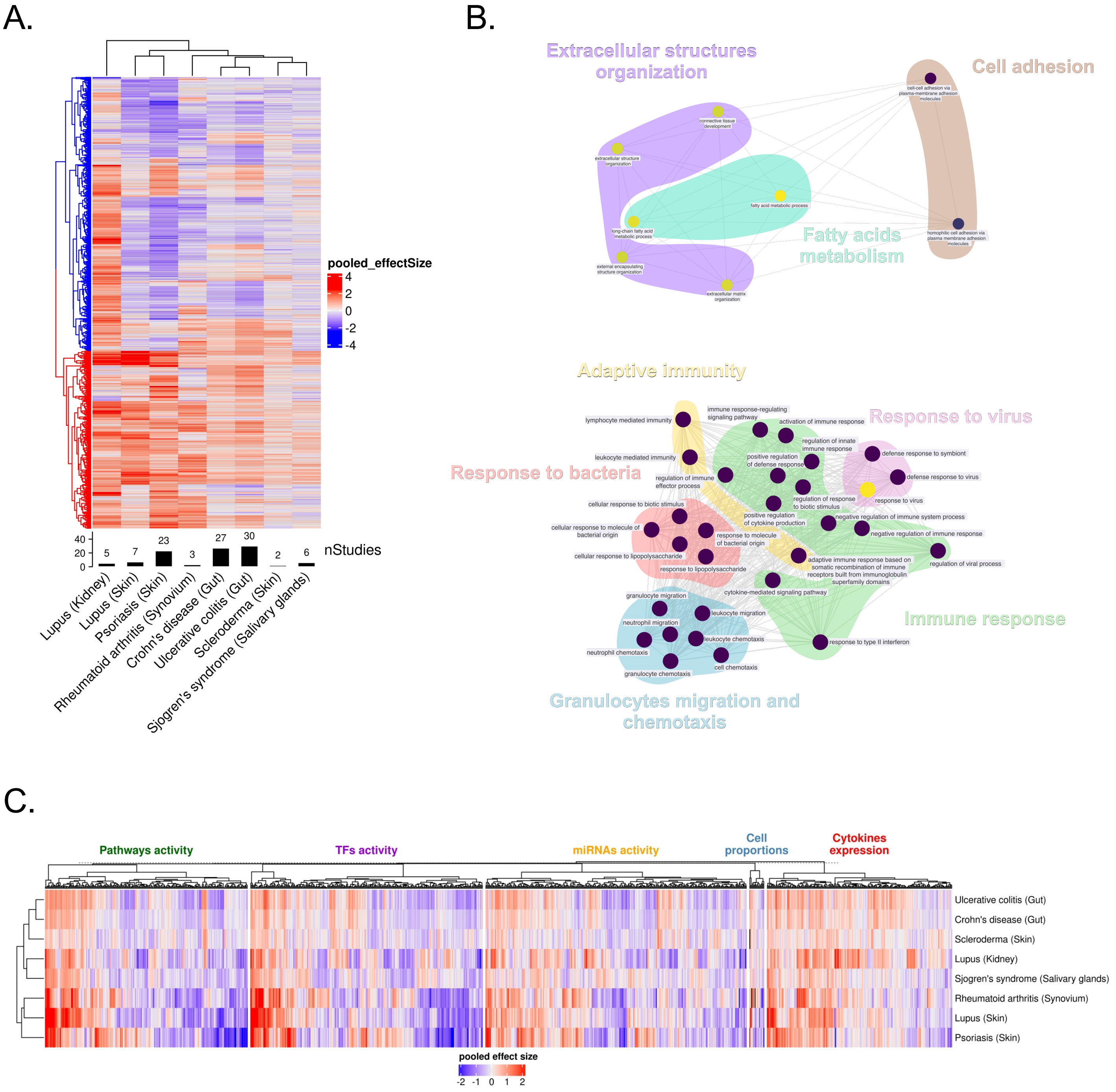

**Supplementary Figure 4.**
**(A)** Gene expression meta-analysis on solid tissue samples. Rows of the heatmap represent genes; only genes with a significant effect size in at least one autoimmune disease are shown. Color indicates the pooled effect size, with red representing upregulation and blue representing downregulation. Hierarchical clustering identifies two gene clusters: one with genes mostly upregulated across diseases (red cluster) and one with genes mostly downregulated (blue cluster). **(B)** Enrichment analysis on the upregulated gene cluster shows significant pathways related to innate and adaptive immunity and viral response (bottom), while the downregulated gene cluster is enriched in tissue structure pathways (top). **(C)** Landscape of dysregulated features in tissue across diseases. Columns represent features (e.g., cell proportions, pathway activities, miRNA activities, transcription factor activities, and cytokine expression) that are significant in at least one disease. Red indicates upregulation, blue indicates downregulation. Rows show disease clustering based on all features.

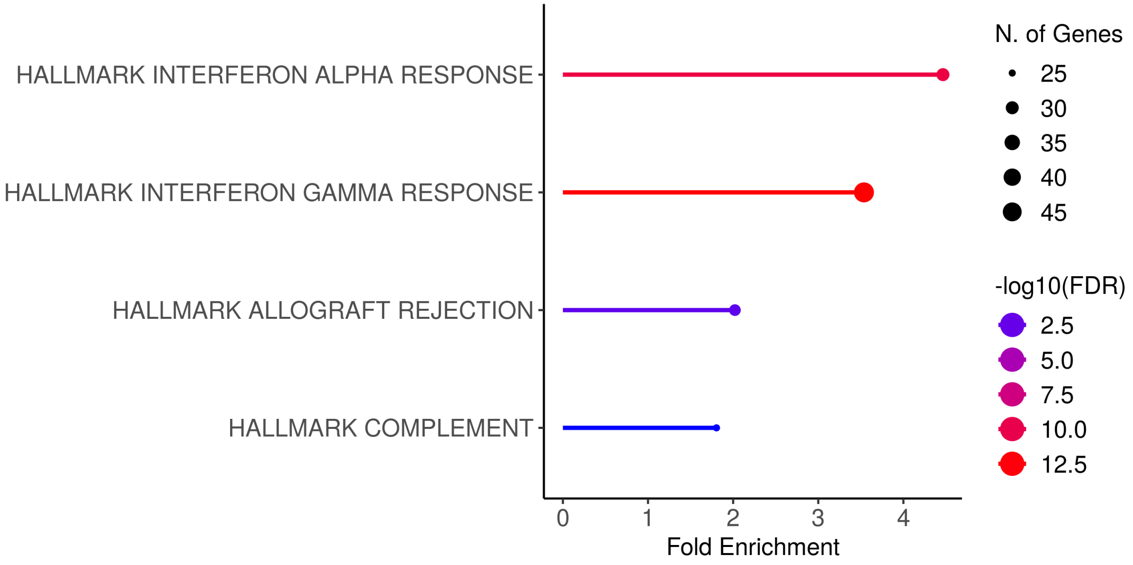


**Supplementary Figure 5.**
Pathway enrichment analysis of genes upregulated in LN: pathway analysis was performed using the ShinyGO 0.82 web server against the MSigDB Hallmark gene sets. Top enriched Hallmark pathways from 1,662 genes upregulated in lupus kidney tissue (FDR<0.05, positive effect size, supported by at least 2 studies) are represented as horizontal lines. Dot size represents the number of genes contributing to each term, while color intensity corresponds to the statistical significance (-$\log_{10}$FDR).

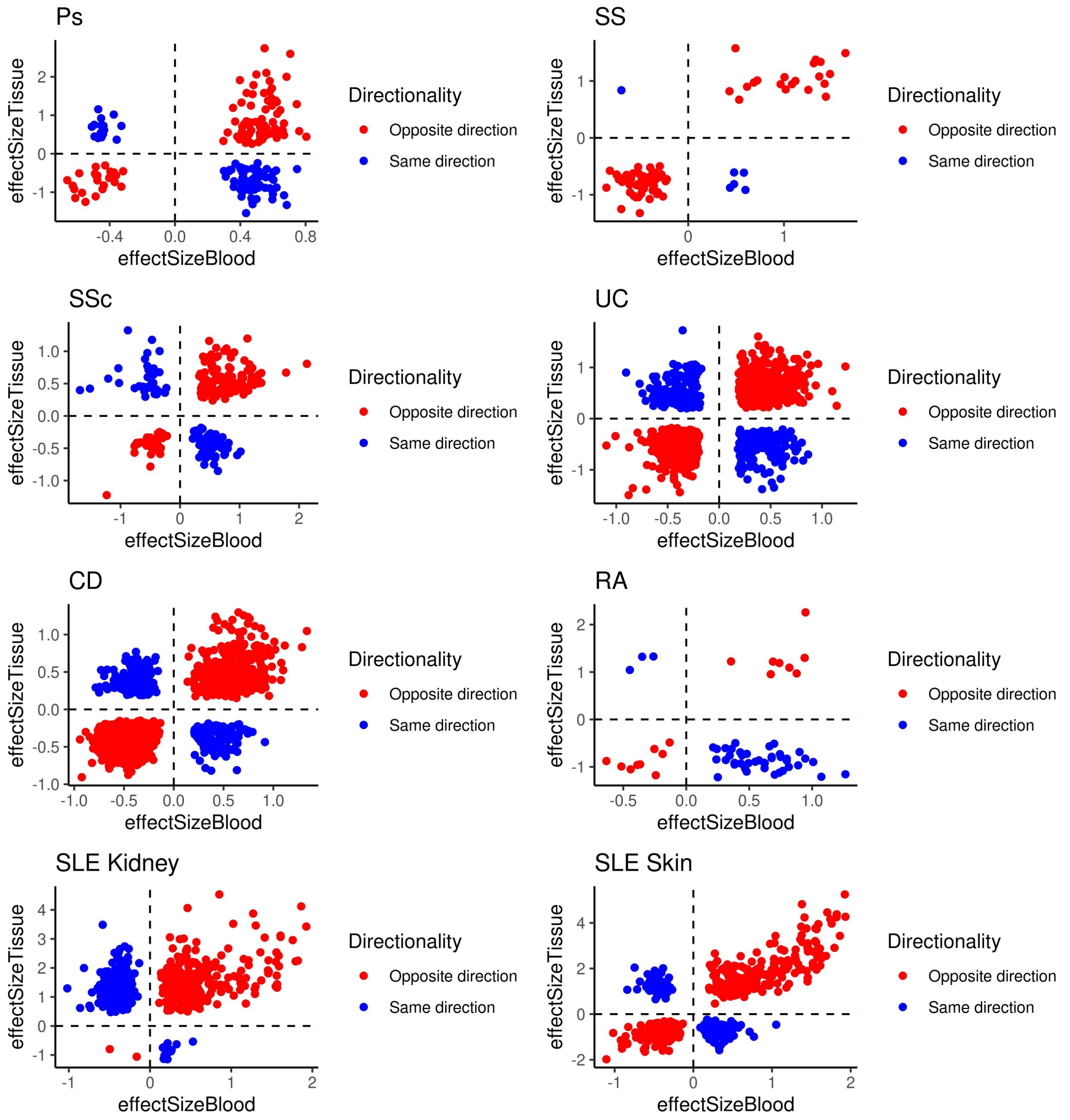


**Supplementary Figure 6.**
Comparison of gene-level effect sizes between blood and tissue samples across autoimmune diseases: Scatterplots display the pooled effect size in tissue (y-axis) versus blood (x-axis) for genes that were significantly differentially expressed (FDR<0.05) in both compartments. Each panel corresponds to a different disease: psoriasis (Ps), Sjögren's syndrome (SS), scleroderma (SSc), ulcerative colitis (UC), Crohn's disease (CD), rheumatoid arthritis (RA), and systemic lupus erythematosus (SLE) in kidney and skin. Genes are colored by directionality: red indicates opposite regulation between compartments, while blue indicates concordant regulation.

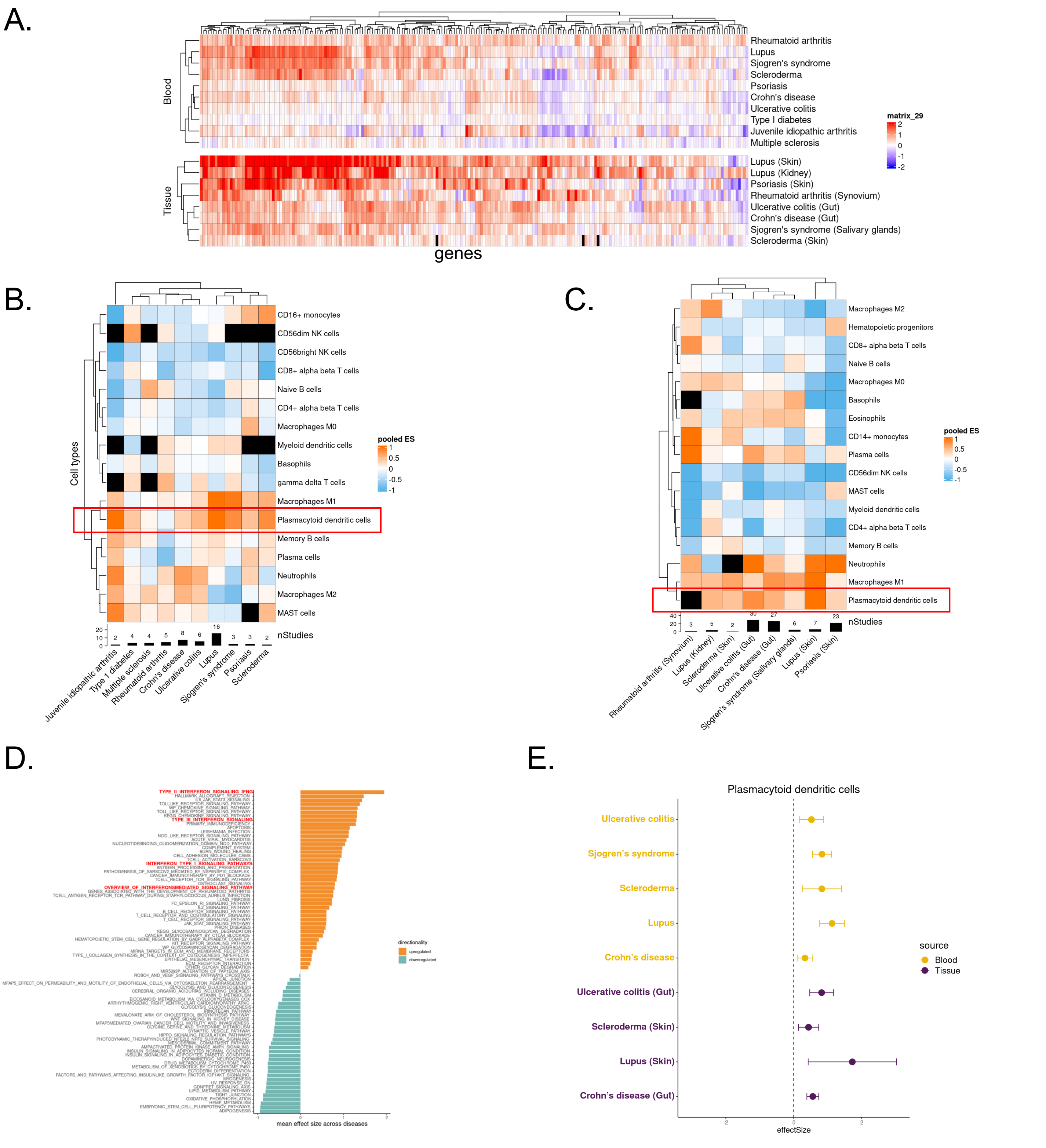


**Supplementary Figure 7.**
**(A)** Interferon α and γ-related genes from hallmark pathways (*HALLMARK_INTERFERON_ALPHA_RESPONSE* and *HALLMARK_INTERFERON_GAMMA_RESPONSE*) effect sizes across diseases. Red indicates upregulation, blue indicates downregulation, shown for both blood and tissue. **(B, C)** Cell type deconvolution meta-analysis results in blood and tissue. Each cell in the heatmap shows the pooled effect size for a specific cell type, with orange representing increased abundance and sky blue representing decreased abundance in disease vs. control. **(D)** Bar plot of mean effect sizes from pathway meta-analysis across diseases shows interferon pathways as the most upregulated across autoimmune diseases. **(E)** Forest plot of pooled effect sizes for pDCs across studies and diseases in both blood (yellow) and tissue (purple), showing only diseases with significant (FDR<0.05) effect sizes.

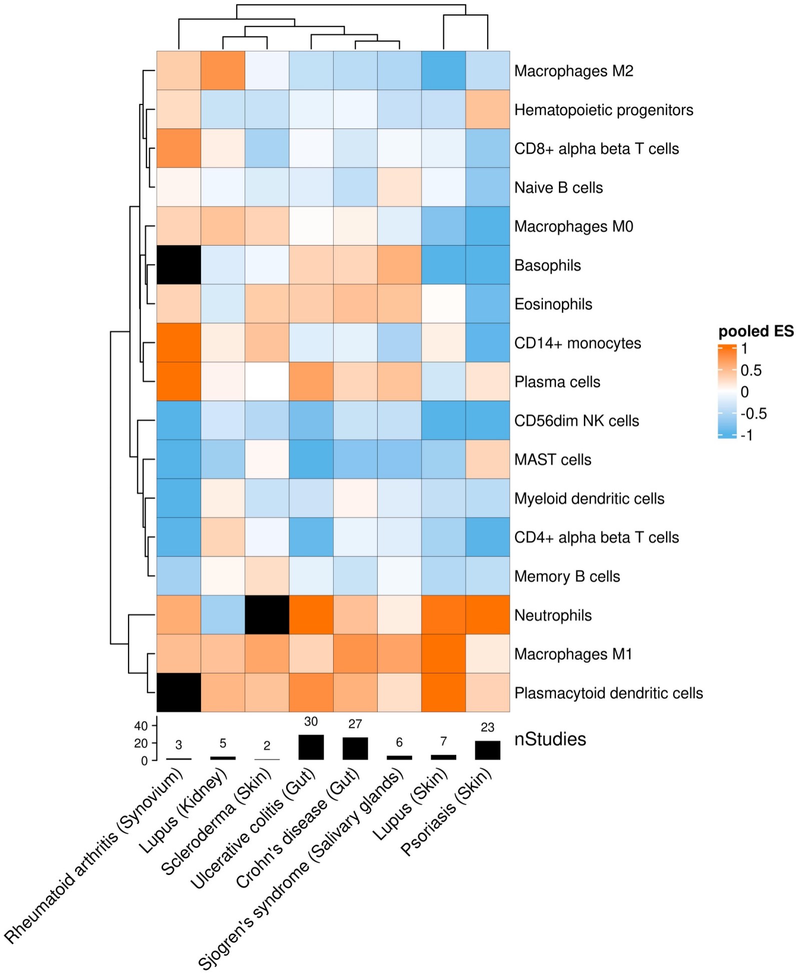


**Supplementary Figure 8.**

Estimated immune cell type proportions in tissue samples across autoimmune diseases. Cell fractions were inferred using computational deconvolution applied to transcriptomic data from affected tissues, and meta-analysis was used to compute pooled effect sizes relative to controls. The heatmap displays differential abundance for 20 immune cell populations across disease-tissue contexts. Black squares indicate the number of contributing studies per condition.

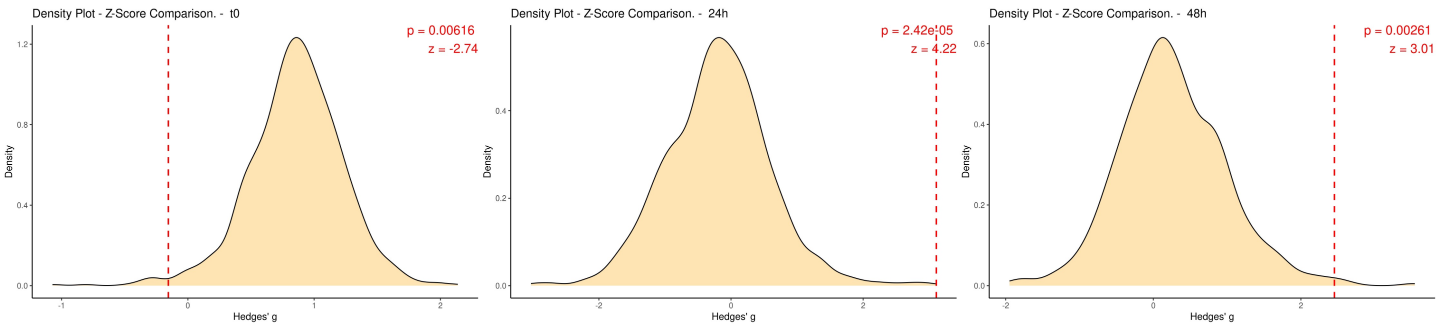


**Supplementary Figure 9.**
Permutation analysis validating the robustness of the IFN module in the perturbation dataset across three time points (0h, 24h, 48h): for each time point, we generated 1,000 random modules (matched in size to the IFN module, 49 features) by randomly sampling features from the remaining 14 modules. We then computed effect sizes (Hedges' g) for both the IFN and randomized modules. Distributions show the density of effect sizes from random modules, while the vertical red dashed lines represent the observed IFN module effect sizes. Z-scores and corresponding p-values were calculated to test whether the IFN module's effect size significantly exceeded random expectation.

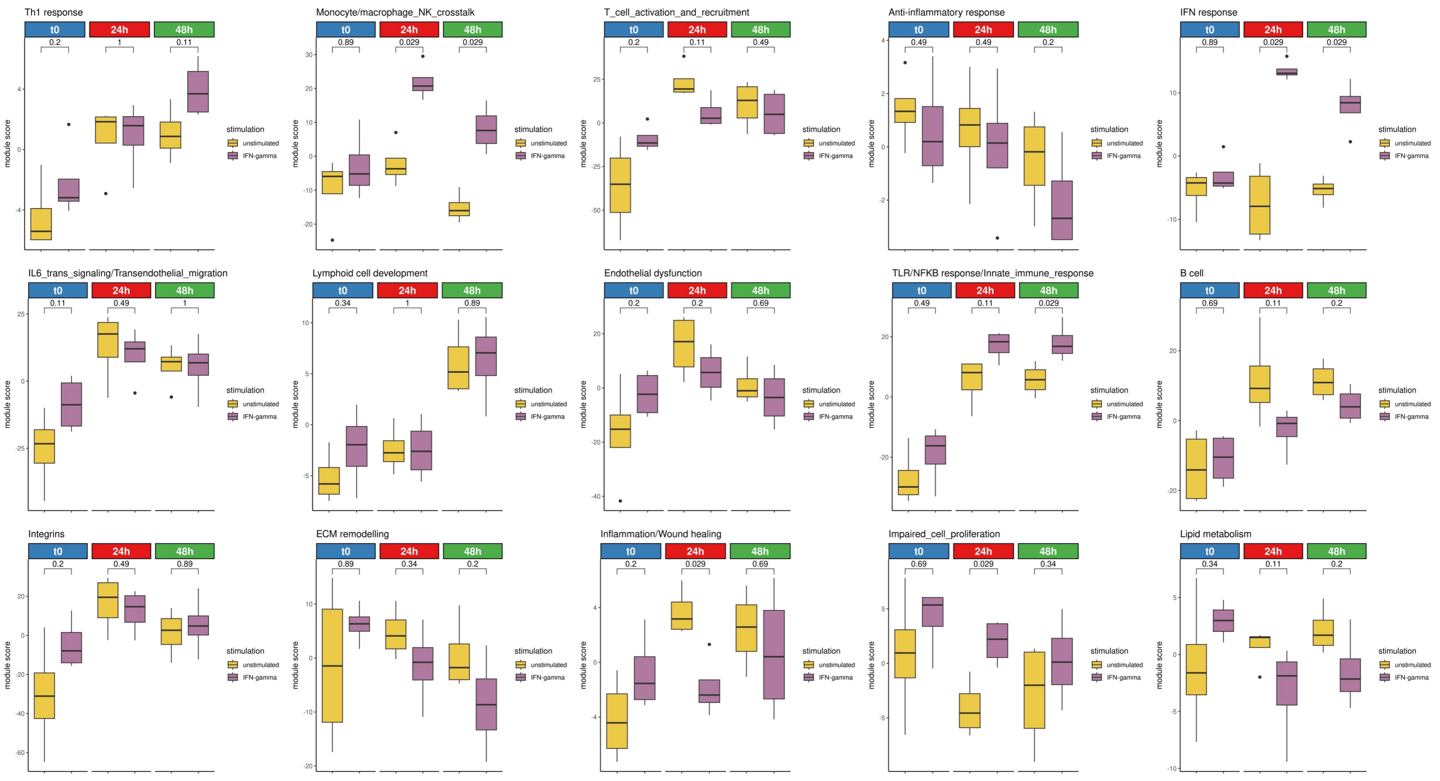


**Supplementary Figure 10.**

Immune module responses to IFN-γ stimulation in whole blood. Module scores are shown for each of the 15 immune modules across three time points: baseline (t0), 24h, and 48h post-stimulation. Statistical comparisons between time-points were performed using the Wilcoxon-test. While the IFN response module shows a strong and specific increase in activity at 24h ($p=0.057$) and 48h ($p=0.029$), most other modules remain unchanged. A few, such as TLR/NFκB_response/innate_immune_response ($p=0.029$), show modest shifts at 48h. The limited responsiveness of the remaining modules underscores the specificity of the IFN module in capturing transcriptional effects of IFN-γ stimulation in this experimental context.

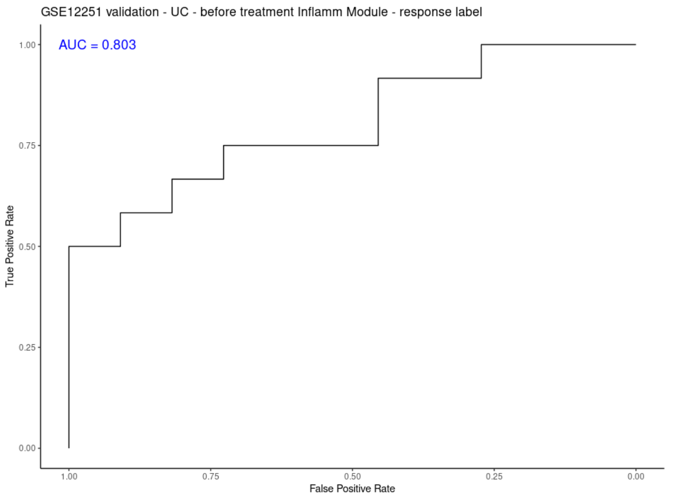


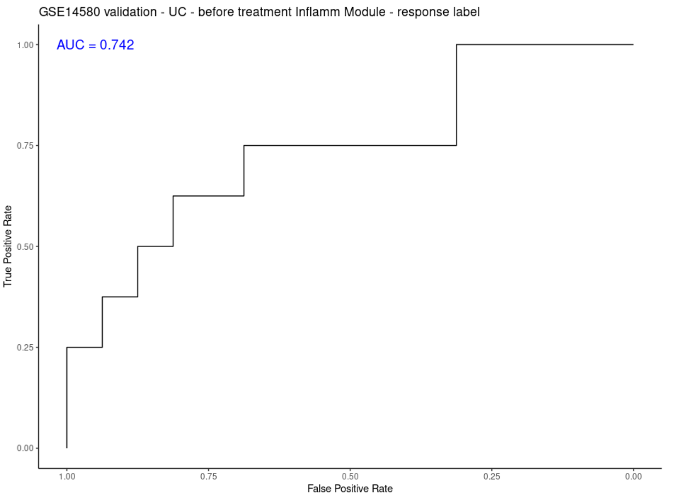


**Supplementary Figure 11.**
Receiver operating characteristic (ROC) curves evaluating the predictive performance of the Inflammation/Wound_healing module in two independent ulcerative colitis (UC) cohorts. Baseline module activity scores were used to distinguish responders from non-responders to infliximab treatment. Classification accuracy was quantified using the Area under the Curve (AUC) metric.
